# Structure and function of the bacterial protein toxin phenomycin

**DOI:** 10.1101/847772

**Authors:** Bente K. Hansen, Camilla K. Larsen, Jacob T. Nielsen, Esben B. Svenningsen, Lan B. Van, Kristian M. Jacobsen, Morten Bjerring, Rasmus K. Flygaard, Lasse B. Jenner, Lene N. Nejsum, Ditlev E. Brodersen, Frans A. A. Mulder, Thomas Tørring, Thomas B. Poulsen

**Affiliations:** Department of Chemistry, Aarhus University, Langelandsgade 140, DK-8000, Aarhus C, Denmark; Department of Engineering, Aarhus University, Gustav Wieds Vej 10, DK-8000, Aarhus C, Denmark; Interdisciplinary nanoscience center (iNANO), Aarhus University, Gustav Wieds Vej 14, DK-8000, Aarhus C, Denmark; Department of Molecular Biology and Genetics, Aarhus University, Gustav Wieds Vej 10, DK-8000, Aarhus C, Denmark; Department of Integrated Structural Biology, Institute of Genetics and Molecular and Cellular Biology, CNRS UMR710, INSERM U964, University of Strasbourg, Strasbourg, 67000, France; Department of Clinical Medicine, Aarhus University, Palle Juul-Jensens Boulevard 82, DK-8200, Aarhus N, Denmark

**Keywords:** Ribosome, protein synthesis inhibitor, natural product, mini-protein, cell-penetrating peptide, protein toxin, NMR structure

## Abstract

Phenomycin is a bacterial mini-protein of 89 amino acids discovered more than 50 years ago with toxicity in the nanomolar regime towards mammalian cells. The protein inhibits the function of the eukaryotic ribosome in cell free systems and appears to target translation initiation. Several fundamental questions concerning the cellular activity of phenomycin have however remained unanswered. In this paper, we have used morphological profiling to show that direct inhibition of translation underlies the toxicity of phenomycin in cells. We have performed studies of the cellular uptake mechanism of phenomycin, showing that endosomal escape is the toxicity-limiting step, and we have solved a solution phase high-resolution structure of the protein using NMR spectroscopy. Through bioinformatic as well as functional comparisons between phenomycin and two homologs, we have identified a peptide segment, which constitutes one of two loops in the structure, that is critical for the toxicity of phenomycin.

## Introduction

The 1950-70s witnessed the discovery of a multitude of complex natural products. The scientific literature from this period is therefore rich with potentially valuable compounds that still to this day have evaded detailed exploration for no apparent reason besides the challenges of navigating often incomplete chemical structures and – occasionally – ambiguous biological activities. Several recent examples have underscored the value of re-investigating such overlooked natural products.^1,2,3,4^ We have a strong interest in structurally unusual peptidic natural products,^5,6,7^ and therefore took notice of a series of reports on the compound phenomycin, which was first isolated in 1967 from *Streptomyces fervens* MA 564-Cl.^8^ Phenomycin was found to possess antitumor activity in several different mouse models^9^ but displayed no antibacterial activity. The peptidic nature of phenomycin was recognized early^8^ but the complete amino acid sequence was not assigned until 1991 showing that phenomycin is a 89 amino acid protein.^10^ Early studies of the mode of action found that phenomycin could attenuate protein synthesis when added to HeLa S3 cells.^11^ The inhibitory effect, 50% inhibition of [^3^H]-Leucine incorporation at app. 10 μM phenomycin, was found to manifest itself slowly, demanding pre-incubation of the cells for 24 hours before addition of the radioactive tracer. Under the same conditions, the protein did not perturb DNA transcription or replication measured using [^3^H]-uridine or [^3^H]-thymidine incorporation, respectively. In a set of nearly parallel studies, another small protein – enomycin – which was isolated from *Streptomyces mauvecolor* and shares 84% sequence identity (and 89% sequence similarity) with phenomycin was found to have very similar properties.^12,13^ Later studies revealed that phenomycin and enomycin are encoded as precursor protein expressed with an N-terminal leader sequence followed by the previously identified proteins.^14,15^ In cell-free preparations, both compounds were found to inhibit translation immediately and sedimentation studies showed an ability to directly bind mammalian ribosomes *in vitro*.^16^ Both compounds could partially interfere with the binding of aminoacyl-tRNAs to ribosomes^11,16^ and appeared to affect initiation more significantly than elongation,^17,18^ however, a detailed mechanistic understanding of the inhibitory effects remains incomplete. We were intrigued by the potential inhibitory effect of phenomycin on translation and the outstanding questions that surround this small protein. For instance, the slow acting nature of phenomycin (and enomycin) in cells suggests that uptake is the limiting factor for toxicity, but the underlying mechanism was never investigated. This is an interesting question, given the strong contemporary interest in cellular uptake mechanism for large peptides/proteins.^19,20^ Compared to known ribosome-inactivating protein (RIP) toxins, such as shiga-toxin or saporin, that work through catalytic modification of the ribosome,^21^ phenomycin is smaller and also does not share significant sequence similarity. In addition, no structures are available for phenomycin (or enomycin). Beyond the RIP toxins, peptide-based inhibitors of the eukaryotic ribosome are in fact quite scarce whereas many smaller peptides are known to bind and inhibit the prokaryotic ribosome.^22,23^ In this article, we report studies of the cellular uptake pathway of phenomycin demonstrating that the protein can access late endosomes. Furthermore, we provide new data that strongly support ribosomal targeting as the toxicity-mediating effect of phenomycin in human cells. Finally, we provide a high-resolution structure of the protein and we identify peptide segments required for toxicity.

## Results and Discussion

### Phenomycin toxicity in mammalian cells is due to inhibition of protein synthesis

We expressed phenomycin in *E. coli* with a cleavable N-terminal His_6_-TEV-tag and used Ni-NTA affinity chromatography to perform initial purification. Following TEV protease-cleavage and removal of the His_6_-tag, we purified the protein to homogeneity by reverse-phase HPLC. The recombinant protein produced (referred to as phenomycin throughout the paper) contains four extra N-terminal amino acids (Gly-Ala-Met-Ala) to facilitate chemical tagging (Table S1, Supplementary information). After lyophilization, phenomycin is readily solubilized in phosphate buffer and retains activity which indicates high structural integrity and/or a strong ability to re-fold. Initially, we investigated the effect of phenomycin on cell viability in MCF7 cells at different treatment timepoints and we observed a significant change (25-fold) in IC_50_-value between 24 h and 48 h and no further potentiation at 72 h (Figure 1a). This slow onset of the growth inhibitory effect was noted also in earlier studies and was hypothesized to be the result of slow penetration into cells.^10^ Next, we conducted viability tests in a range of cell lines to assess the generality of phenomycins growth inhibitory activity (Figure 1b). The protein was active in all cell lines tested and showed >10-fold differential activity across the set, with MDA-MB-231 cells (breast cancer) being most sensitive (IC_50_ = 0.26 ± 0.2 μM) and murine AML12 liver cells least sensitive (IC_50_ = 3.8 ± 0.2 μM). In comparison, the tripeptide MG-132, which is an inhibitor of the 20S proteasome, was equally potent in all cell lines (<2-fold variation in IC_50_-value, Figure S1).

**Figure 1.**
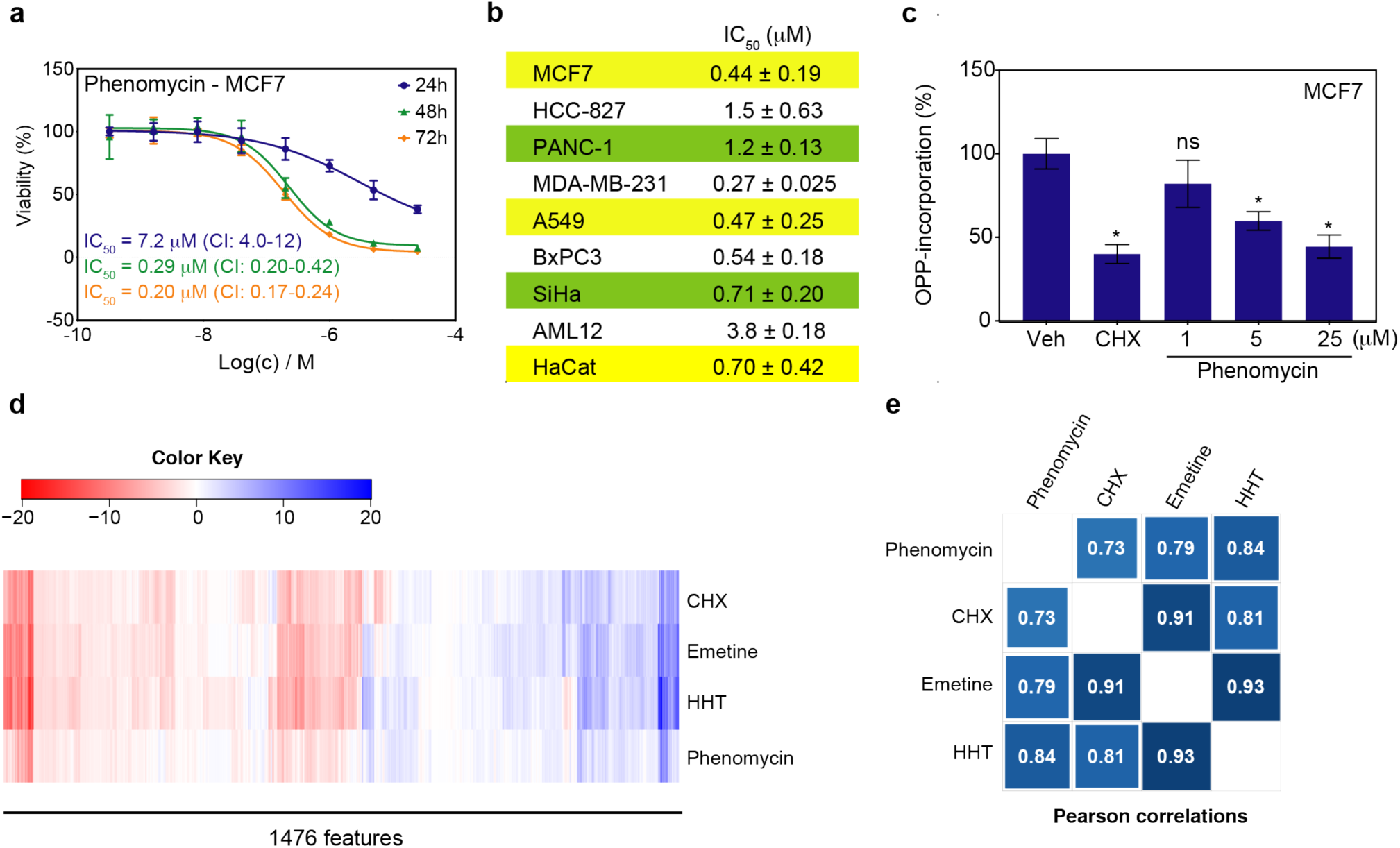
Phenomycin is toxic to mammalian cells and cluster mechanistically with inhibitors of translation. (a) Time-dependent inhibition of cell viability by phenomycin in MCF7 cells. Data points are mean ± s.d. (*N*=3). CI = 95% confidence interval. (b) IC_50_-values of phenomycin in various cell lines (72 h). Values are mean ± s.d. of triplicate (HCC-827, MCF7, A549) or duplicate (PANC-1, MDA-MB-231, BxPC3, SiHa, AML12, HaCat) biological repeats, each of which was performed in triplicate. (c) Relative protein synthesis in MCF7 cells measured by OPP (O-propargyl-puromycin) incorporation for 30 min following initial compound treatment. Cycloheximide (CHX) treatment was at 10 μM for 30 min. Phenomycin treatment was for 12 hours. Vehicle (Veh) was PBS. Data are mean ± s.d. (*N*=3) from a representative experiment performed three times with consistent outcome. ns = non-significant. * *P* < 0.005 relative to vehicle-treated cells (Students t-test, unpaired). (d) Morphological profiles of phenomycin and three distinct inhibitors of translation (5 µM). U2OS cells are treated for 24 hours in four replicates which are averaged to one fingerprint. HHT = homoharringtonine. (e) Pearson correlations of morphological profiles for phenomycin and translation inhibitors. See also Figure S1 and Table S2.

To conclusively connect the cellular mechanism of toxicity to inhibition of translation, we performed two complementary experiments: First, we directly assessed the ability of phenomycin to inhibit protein synthesis in MCF7 cells using a puromycin-based assay to label newly synthesized peptides and found dose-dependent inhibition (Figure 1c). However, compared to cycloheximide (CHX), which strongly inhibits puromycin-incorporation after only 30 minutes, treatment with 5 μM phenomycin (10 x IC_50_ at 72 h) required up to 12 hours to show significant inhibition. This slow-acting effect is similar to the toxicity profile and suggest that the two effects are coupled. In a more decisive set of experiments, we subjected phenomycin to morphological profiling (24 hours) in U2OS cells.^24,25,26^ Morphological profiling is a computational inference technique^27^ based on fluorescence imaging that can be used to mechanistically classify bioactive compounds through their effects on the cellular ultrastructure, imaged using six different organelle-targeted fluorophores (Figure 1d and Table S2). We used three inhibitors of protein synthesis - CHX, emetine, and homoharringtonine (HHT) – as direct experimental controls. Despite having distinct inhibitory effects on translation,^28^ the latter compounds have strongly correlated bioactivity profiles and constitute a distinct cluster in our current database (Figure S1). Likewise, phenomycin was found to afford a bioactivity profile with high correlations (Pearson correlation coefficients 0.73-0.84) to these compounds which is indicative for overall mechanistic similarity (Figure 1d,e). Collectively, these experiments strongly suggest that inhibition of ribosomal protein synthesis is directly responsible for the toxicity of phenomycin in mammalian cells.

### Phenomycin is present in cellular endosomes and toxicity is enhanced by protein delivery agents

The type 2 RIP toxins, such as ricin and shiga-toxin, utilize elaborate cell-entry mechanisms, facilitated by their B-subunit(s), to deliver the catalytic A-subunit to the cellular cytoplasm where it directly depurinates specific nucleotides in the 28S rRNA and irreversibly blocks translation elongation.^21,29^ Type 1 RIPs, e.g. saporin, consist of only the 30 kDa A-subunit and are therefore slower to penetrate into cells. By comparison, the modest size (9.5 kDa) of phenomycin in combination with its polycationic nature (10 lysines, 4 arginines, Table S1) initially suggested to us that an uptake mechanism reminiscent of cell penetrating peptides (CPPs),^30^ such as TAT or polyarginines, might be involved. To enable studies of the cellular uptake of phenomycin, we engineered a cysteine-residue into the N-terminal section of the protein (M3C phenomycin, Table S1) for selective conjugation. Alkylation with an iodoacetamide-derivative having a terminal alkyne (IA-alkyne) was used to install a biorthogonal handle to the protein (Figure 2a). Using a copper(I)-catalyzed click reaction^31,32,33^ we could then readily conjugate sulfo-Cy5-azide or FAM-azide to generate fluorophore-labelled phenomycins for uptake studies in live cells (Figure 2a and Figure S2). We also generated a biotin-labelled conjugate through the same overall workflow and we confirmed that all phenomycin conjugates maintained toxicity in mammalian cells (Figure S2). When we performed uptake studies of sulfo-Cy5-phenomycin in live MCF7 cells using confocal laser scanning microscopy, we observed the presence of the labelled protein in vesicular structures. Transient transfection of the cells with GFP-Rab7, a marker for late endosomes, followed by sulfo-Cy5-phenomycin treatment led to significant co-localization (Figure 2b) which suggests that phenomycin accumulates in endosomes following initial uptake via endocytosis. In accord with the latter, we found that the uptake of phenomycin could be blocked at 4 ^o^C similar to transferrin (Figure S2). Through time course studies, we observed enhanced vesicular uptake from 1 h to 24 h with 1 μM of sulfo-Cy5-phenomycin but no clear indication of cytoplasmic distribution of the labelled protein. At a higher concentration (5 μM) significant toxicity was observed at the 24 h timepoint. To further investigate whether cellular uptake constituted the toxicity-limiting factor for phenomycin, we used BioPORTER to increase uptake efficiency. The BioPORTER reagent can both mediate direct cellular translocation of proteins across the plasma membrane as well as increase endosomal escape (Figure 2c). While the BioPORTER reagent alone did not afford any measurable toxicity (Figure 2d), the combination with phenomycin resulted in a nearly 10-fold increase in potency (Figure 2e). Taken together, these studies demonstrate that phenomycin is efficiently internalized in cells by an energy-dependent process and that access to the cytoplasm – through endosomal escape^34^ – is likely required for toxicity.

**Figure 2.**
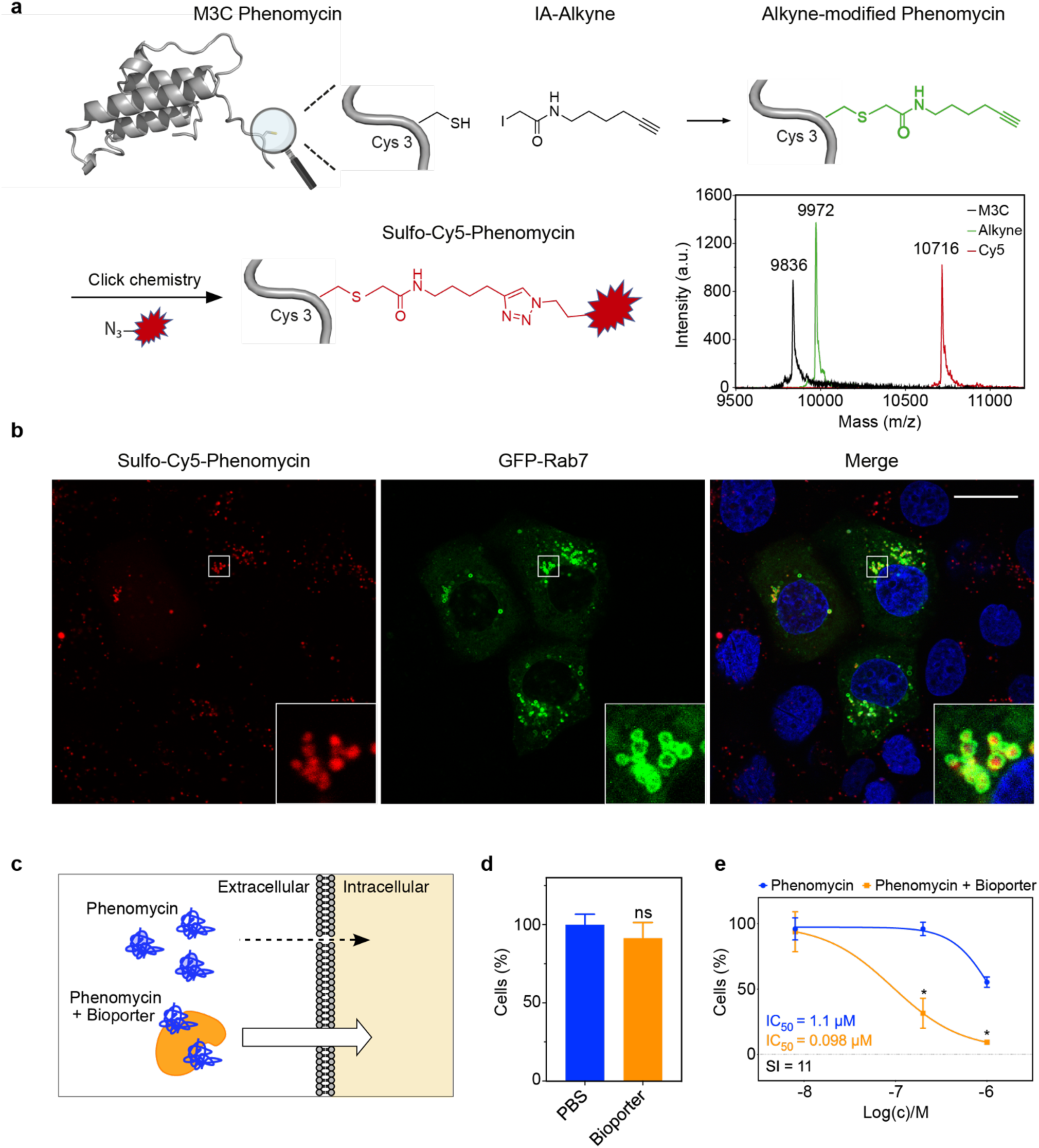
Fluorophore labelled phenomycin is present in late endosomes and toxicity is enhanced by a protein delivery agent. (a) M3C phenomycin is alkylated by IA-alkyne in the presence of TCEP and the resulting conjugate can be purified by preparative reverse phase HPLC. Copper-catalyzed azide-alkyne cycloaddition enables conjugation with sulfo-Cy5 followed by preparative reverse phase HPLC purification. MALDI-TOF mass spectrometry (uncalibrated) of the different conjugates is shown. Calculated masses: M3C: 9826.08; Alkyne: 9963.16; Cy5: 10726.14 (b) Colocalization study of sulfo-Cy5-labelled phenomycin (red) and transiently expressed GFP-Rab7 (green) as a marker of late endosomes in MCF7 cells reveals phenomycin to be present in Rab7-positive vesicles. Cell nuclei are stained by Hoechst 33342 (blue). Sulfo-Cy5-phenomycin was used at 5 μM and treatment time was 12 h. Scale bar is 20 µm. 8-bit images with display range: 0-190. (c) The BioPORTER reagent can enhance cellular uptake of proteins through different mechanisms. (d) The BioPORTER reagent alone is not cytotoxic to MCF7 cells after 72 h treatment. Data points are mean ± s.d. (*N* = 3), ns = non-significant relative to PBS-treated cells. (e) Co-treatment of MCF cells (72h) with phenomycin-BioPORTER enhances toxicity relative to phenomycin alone. Curves are from a representative experiment repeated twice with consistent outcome. Data points are mean ± s.d. (*N* = 3). * *P* < 0.005 relative to phenomycin-only treated cells. SI = selectivity index. See also Figure S2.

### Solution structure of phenomycin determined by NMR spectroscopy

Given its interesting functional properties, we next sought to determine a solution structure of phenomycin by NMR spectroscopy. To this end, we expressed phenomycin in minimal medium to introduce ^13^C and ^15^N-isotope labels. After HPLC purification the lyophilized protein was dissolved in phosphate buffer containing 5% D_2_O and NMR experiments were performed. The structure of phenomycin (Figure 3) was calculated using 1901 distance constraints of which 543 were long range. The maximum distance and torsion angle restraint violation have low values of 0.16 Å and 3.05° (on average), respectively. The structure ensemble has high precision with a rmsd of 0.30 and 0.76 Å for backbone atoms and heavy atoms, respectively, for the structured part (residues 5-93). All restraint and structure calculation statistics are provided in Table S3. The structure is shown in Figure 3, and a schematic cartoon of the two-dimensional structure is shown in Figure S3. Phenomycin consists of three α-helixes, *H1* (A11-A24), *H2* (V43-D61) and *H3* (I75-M88) that form a three-helix bundle in which *H1* and *H2* are oriented parallel to each other and *H3* is anti-parallel (Figure 3a). The helical bundle is capped by a small anti-parallel β-sheet comprising two β-strands, β1 (T8-K10) and β2 (D40-P42). The helices are connected by two well-structured loops. Loop 1 (G25-P42, Figure 3e) which contains a 3_10_-helix (A29-K31) with H-bonds between T28 and K31 as well as T28 and S32 and an extended linker segment (H33-T39) that connects to β2 and enables the parallel orientation of helices *H1* and *H2*. This linker is anchored with I35 that makes several hydrophobic contacts to residues A11, Y14, V41 and Y45 in the hydrophobic core. Loop 2 (K62-S74, Figure 3f) also starts with a 3_10_-helix (A66-D68) with H-bonds between P65 and D68 as well as A66 and K69 and ends with a small β-hairpin connecting strands β3 (K69-K70) and β4 (M73-S74). The fold of the loop is stabilized by side chain hydrogen bonds between the positively charged sidechain amino group of K69 and both L64/D61 main chain carbonyl groups and is close enough to form salt-bridges to the negatively charged side-chains of D61 and D72 although such conformations could not be confirmed in all NMR models in the structure ensemble. The C-terminal residues (I90-W93) are relatively structured with W93 packing against the hydrophobic core and D91 and T92 having side-chain H-bonds to N5. The extra residues 1-4 added artificially to the N-terminus of the native phenomycin sequence are unstructured, having no specific long-range interactions with the rest of the structure.

**Figure 3.**
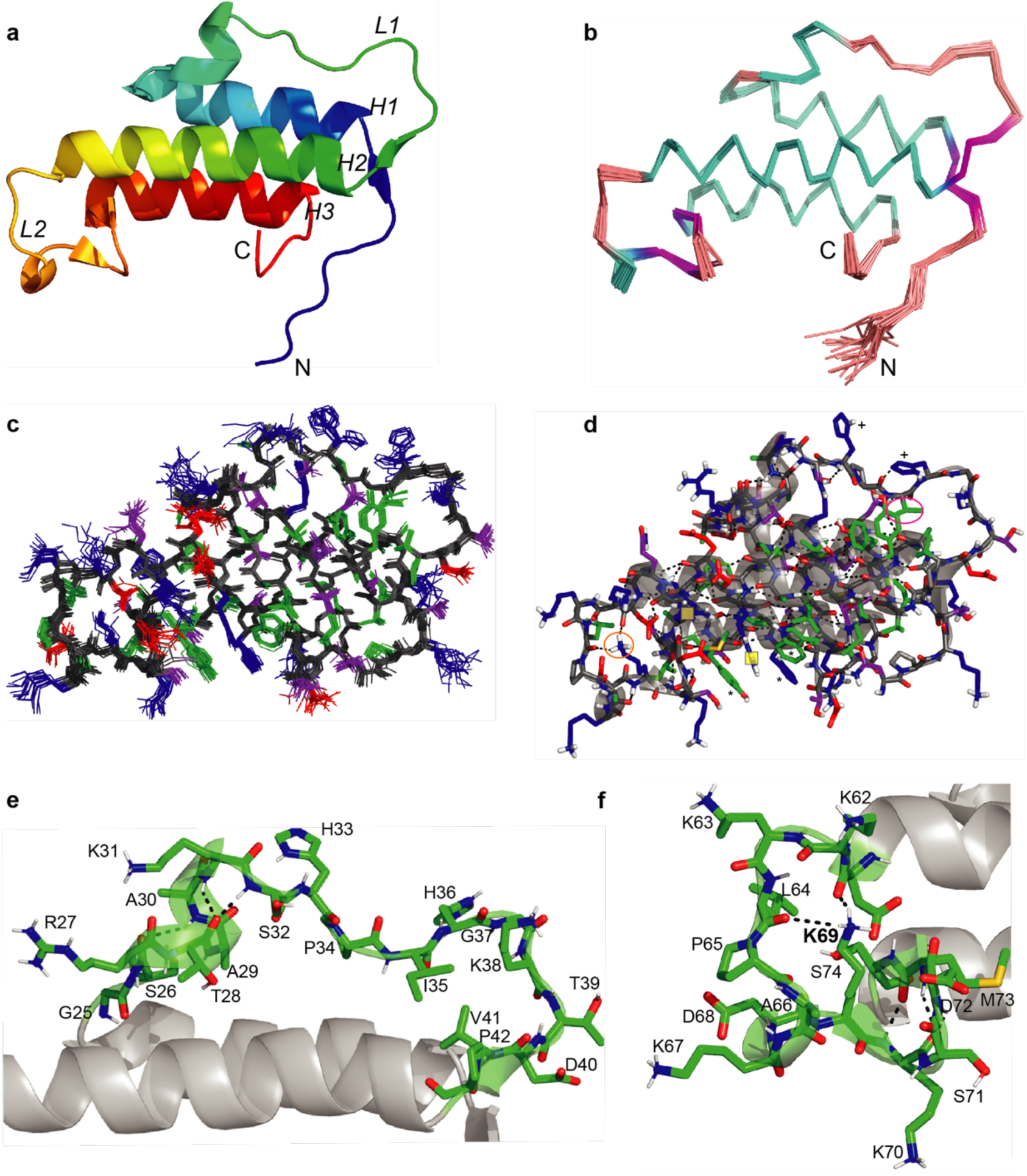
Solution structure of phenomycin. Only residues 5-93 are represented in panels c-f. (a) Cartoon representation using rainbow colors from N (blue) to C (red) terminus. (b) Ensemble ribbon representation colored according to secondary structure (helix: cyan, sheet: magenta. (c) Line representation showing the C, N, and O atoms of all NMR models with main-chain atoms in grey and using residue class-specific coloring for side chain atoms: aliphatic (green), Gly/Pro (yellow/grey), positively charged incl. His (blue), and negatively charged (red). (d) Stick-representation of the lowest energy model using the same coloring scheme for carbon atoms as in (c), showing N/O/H-atoms in blue/red/white (only showing polar hydrogen bonds), and displaying secondary structure elements with transparent grey cartoons and hydrogen bonds with black dashes. Specific side chains mentioned in the text are highlighted: The NH_3_^+^ group of K69 (orange ellipse), I35 (magenta ellipse), R54/R58 guanidinium groups (yellow boxes), H33/H36 (“+”), aromatic Y78/H82/W93 (“*”). (e) Close-up showing loop 1 in a stick representation using atomic colors as in (c) and carbon atoms in green and residue numbers annotated. (f) Close-up showing loop 2 with representation as in (e). See also Figure S3.

Overall, the phenomycin structure presents a tightly packed hydrophobic core and a surface that is covered with polar and charged residues, where, in particular, more than 10 positively charged side-chains are exposed (Figure 3c,d). This distribution of amino acids is facilitated by the orientation of the three amphipathic helices that orient the hydrophobic sidechains towards the hydrophobic core while locating the hydrophilic and charged residues on the surface (Figure S3). The hydrophobic core is further stabilized by aliphatic and aromatic residues from the loops and strands with I9, V43, I35, M73, and L64 forming a closely packed interior (Figure S3). In contrast, loop 2 contains 5 lysine residues and is close to the C-terminal end of helix 2 that contains two positively charged residues, R54 and R58.

### Phenomycin homologs carotomycin and aquimarinamycin enter endosomes but lack cellular toxicity

Aiming to identify the structural elements in phenomycin that are required for biological activity, we searched for similar protein sequences in the hope that conserved residues would correspond to residues essential for the observed activity. A BLAST search yielded a large group of very similar sequences from other actinobacteria, including the before-mentioned enomycin, and, curiously, unlike phenomycin and enomycin, most of the hypothetical proteins found do not appear at first sight to be encoded with N-terminal leader peptides. We chose to focus on the mature phenomycin and an alignment of the predicted peptides from other actinobacteria can be seen in Figure S4. There is high sequence similarity throughout most of the sequence with the exception of the part annotated as loop 2 (see Figure S4). The more dissimilar alignment in this part does not alter the fact that all members contain a high frequency of basic amino acids, lysine and arginine (avg. 5, s.d. 1). Looking past these close homologs, a predicted protein from *Pectobacterium carotovorum* - a Gram negative plant pathogen – is conspicuous. Williamson and co-workers identified a transposon mutant where the phenomycin-like gene was up-regulated together with a nonribosomal peptide synthetase-polyketide synthase (NRPS-PKS) hybrid pathway producing an orange pigment of unknown structure.^35^ It was demonstrated that the transposon mutant was hypervirulent in potatoes and recognizing the homology to phenomycin and enomycin, the authors hypothesized a similar bioactivity.^35^ The peptide (48% identical, 65% similar to phenomycin), which we will term carotomycin, has a similar predicted secondary structure to phenomycin, but major differences in the two loop regions - including the absence of the characteristic lysine residues present in loop 2 of phenomycin (Figure 4). Further down the list, we found a predicted protein from *Aquimarina muelleri* that appeared similar, but with the noticeable difference of lacking the entire Loop 2 (see Figure 4). The polypeptide, which we will term aquimarinamycin, shows the same predicted pattern of α helices as phenomycin and carotomycin, but only aligned with the N-terminal part of phenomycin leading up to the second loop. We chose to express both carotomycin and aquimarinamycin to compare their bioactivity, cellular uptake, and ribosome affinity with those of phenomycin.

**Figure 4.**
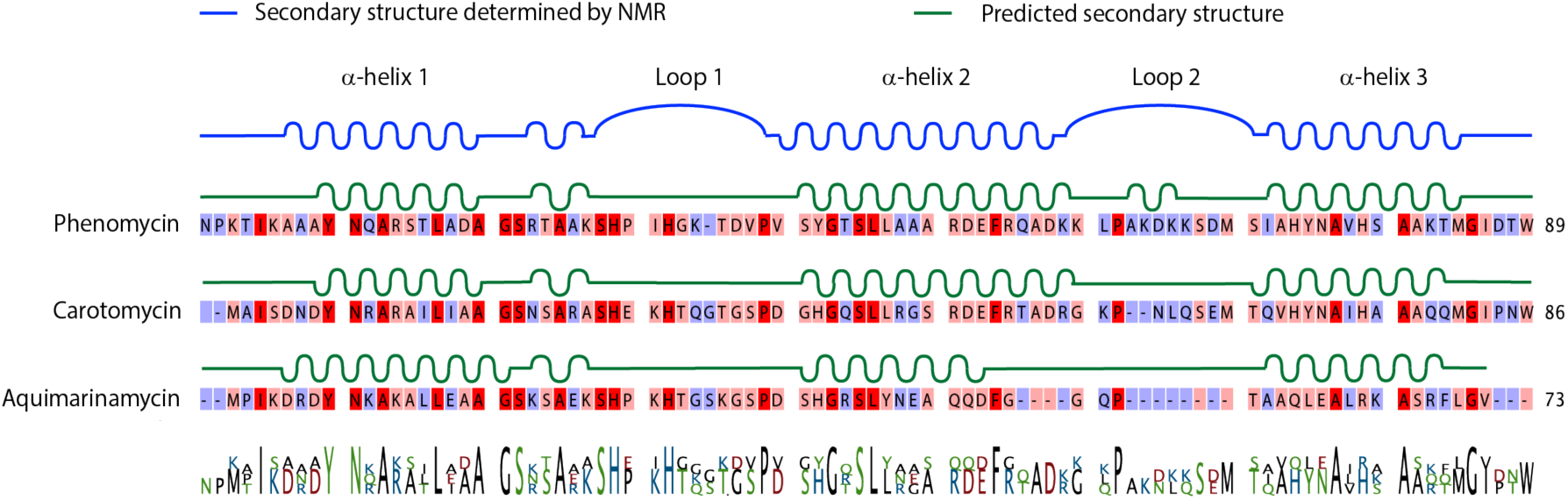
Alignment and secondary structure of phenomycin homologs. The protein sequence of the mature phenomycin is aligned with carotomycin from *Pectobacterium carotovorum* and aquimarinamycin from *Aquimarina muelleri*. One-letter amino acids are color-coded according to similarity (red: 3 of 3, pink: 2 of 3, blue: none). Loop 2 differs most between the two homologs. A cartoon shows the secondary structure determined by NMR for phenomycin (blue) and the predicted secondary structure for all three proteins (green). The NMR determined secondary structure is well in agreement with the predicted structure. The predicted secondary structure of the three peptides appear very similar. The consensus sequence is listed below. The secondary structure was predicted using JPred 4^36^. See also Figure S4.

We inserted the codon-optimized genes into pETM11 in frame with an N-terminal His_6_-TEV-tag similar to the strategy used for phenomycin and followed the same expression and purification protocol. The two polypeptides expressed and purified well, but they showed no activity in MCF7 cells when tested in parallel with phenomycin (Figure 5a). We hypothesized that the loss in growth inhibitory activity could arise from an inability to permeate the cells and/or a decreased affinity towards the ribosome and with respect to the latter, we used biolayer interference (BLI) to investigate the direct interaction with the ribosome. Similar to phenomycin, we prepared cysteine mutants also of carotomycin and aquimarinamycin to enable conjugation to biotin, immobilized the labelled proteins on streptavidin-coated sensor surfaces, and measured association and dissociation curves to 80S ribosomes isolated from HEK293 cells in various concentrations. We found that phenomycin binds very potently with a calculated K_d_ of 23 pM based on a 1:1 binding model (Figure 5b). This data thus provides a quantitative measure on the interaction originally observed by sedimentation analyses.^16^ To investigate differences in species selectivity, we also tested affinity to yeast, plant and bacterial ribosomes. Phenomycin maintained an apparent high affinity for both plant and yeast ribosomes but the traces could not be fitted to a 1:1 binding model with high confidence thus making exact K_d_ determinations difficult (Figure S5). Association to bacterial ribosomes could also be detected but again the interaction did not fit a 1:1 model however, based on the rapid dissociation, the affinity was markedly reduced (Figure S5). In comparison, neither carotomycin nor aquimarinamycin showed any association to any of the ribosomes (Figure S5). We recorded circular dichroism (CD) spectra of all three polypeptides which demonstrated that carotomycin is very similar to phenomycin in terms of secondary structure but displays reduced temperature stability (Figure 5c, Figure S5). In comparison, aquimarinamycin is unstable or intrinsically disordered (Figure 5c, Figure S5). Finally, to compare uptake with phenomycin, we prepared fluorophore labelled versions of both carotomycin and aquimarinamycin - albeit with orthogonal fluorophores (sulfo-Cy5 for carotomycin/aquimarinamycin and FAM for phenomycin) to enable co-localization studies. As is evident from Figure 5d, both proteins enter cells and co-localize with phenomycin in vesicles, suggesting an identical uptake pathway. Collectively, we have found that phenomycin displays picomolar affinity for the human 80S ribosome. Given this degree of potency and the significant similarity between phenomycin and carotomycin, the lack of affinity to any ribosomes observed for the latter is surprising.

**Figure 5.**
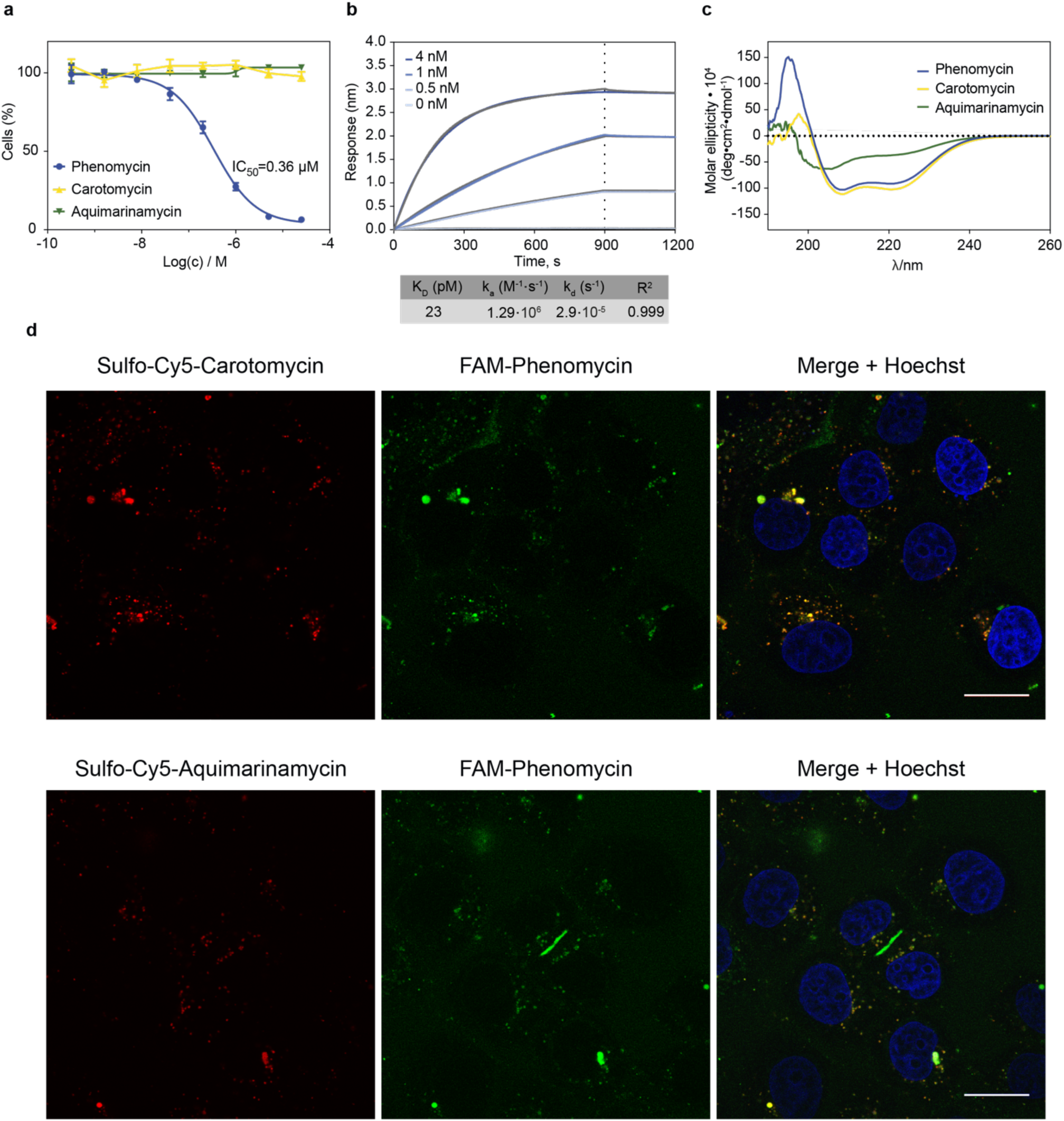
Phenomycin homologs are not cytotoxic, lack ribosome binding activity and co-localize with phenomycin in intracellular vesicles. (a) Measurement of cell viability of the three proteins in MCF7 cells (72 h) demonstrates that only phenomycin is active. Curves are from a representative experiment repeated three times with consistent outcome. Data points are mean ± s.d. (*N* = 3) (b) BLI measurements of phenomycin and human 80S ribosome show that phenomycin binds tightly to the human ribosome. Association/dissociation curves for immobilized phenomycin and human ribosome in varying concentrations (blue-toned lines) and a 1:1 binding fit is shown in grey. Based on the curves calculated binding constants are generated and shown below. The data is from a representative experiment. (c) CD spectra at ambient temperature (25°C). While carotomycin and phenomycin appear very similar in their secondary structure, aquimarinamycin appears disordered. (d) Sulfo-Cy5-conjugates of carotomycin and aquimarinamycin (5 μM) were separately co-incubated with FAM-labelled phenomycin (5 μM) in MCF7 cells for 12 h and the live cells were imaged using confocal microscopy. Cell nuclei are stained by Hoechst 33342 (blue). Scale bar is 20 µm. *Top row:* cellular uptake of a combination of sulfo-Cy5-carotomycin (*left*), FAM-phenomycin (*middle*), and overlay of Hoechst stain, Cy5 and, FAM (*right*). 8-bit images, display ranges: 0-70. *Bottom row:* cellular uptake of a combination of sulfo-Cy5-aquimarinamycin (*left*), FAM-phenomycin (*middle*), and overlay of HOECHST stain, Cy5 and, FAM (*right*). 8-bit images, display range: 1-120 (sulfo-Cy5-carotomycin and Hoechst) and 3-70 (FAM-phenomycin). Both carotomycin and aquimarinamycin co-localize with phenomycin in intracellular vesicles which suggest an identical uptake pathway. Scale bar is 20 µm. See also Figure S5.

### Mutational studies demonstrate that loop 1 of phenomycin is necessary for toxicity

Next, we wanted to analyse the difference in biological activity between phenomycin and carotomycin. Specifically, we decided to focus on the two loop-regions and constructed a series of hybrids between phenomycin and carotomycin to investigate if we could knock-out function (as measured by ability to reduce cell viability) in phenomycin by substituting with the loop sequences from carotomycin or if we could knock-in function by substituting the phenomycin loops into the otherwise inactive carotomycin. With respect to the latter approach, neither of the phenomycin loops could augment carotomycin into measurable cytotoxicity (data not shown). Attempting to knock-out activity of phenomycin, we found that loop 2 is very tolerant of changes. We first mutated the stabilizing lysine-residue of loop 2 to alanine (K69A, see also Figure 3f) but this resulted in no change in activity (Figure S6). In fact, the complete loop 2 of phenomycin could be substituted with the corresponding predicted loop from carotomycin without detrimental effect (Figure S6). In contrast, we found that loop 1 was essential for phenomycin bioactivity as substituting this with the corresponding predicted loop from carotomycin resulted in an inactive hybrid (Figure S6). To get a more detailed understanding of the specific amino acids required for activity, we dissected this loop by exchanging three amino acids at a time with alanine triplets or quartet (Figure 6a). In addition, we created a PI->EK mutant that we saw as a major difference between carotomycin and phenomycin. Interestingly, changes to first part of loop 1 (KSH->AAA or PIH->AAA) resulted in complete loss of activity (IC_50_ > 25 μM, Figure 6b) while other changes to the loop sequence resulted in ca. 3-fold (GKT->AAA) or 15-fold (DVPV->AAAA and PI->EK) loss of activity. CD measurements revealed that mutants KSH, PIH, GKT and DVPV maintained a secondary similar to phenomycin (Figure 6c, Figure S6). The DVPV and PI/EK mutants tended to unfold at lower temperature (Figure S6) indicating decreased stability. Morphological profiling of all mutants and hybrids corroborated these observations: Whereas the KSH and PIH-alanine triplet mutants had no clear profiles, the remaining constructs were active and afforded profiles with strong correlations to phenomycin, which is suggestive of a shared mechanism of action (Figure S6). Collectively, the sequence comparison between carotomycin and phenomycin and the mutational studies indicate that the sequence **KSH**PI**H** appears to represent a critical structural component of phenomycin. A close inspection of the structure of loop 1 (Figure 6a and Figure 3e) shows that K31 and the two histidines (H33 and H36) are all located on the surface of the protein and are flexible.

**Figure 6.**
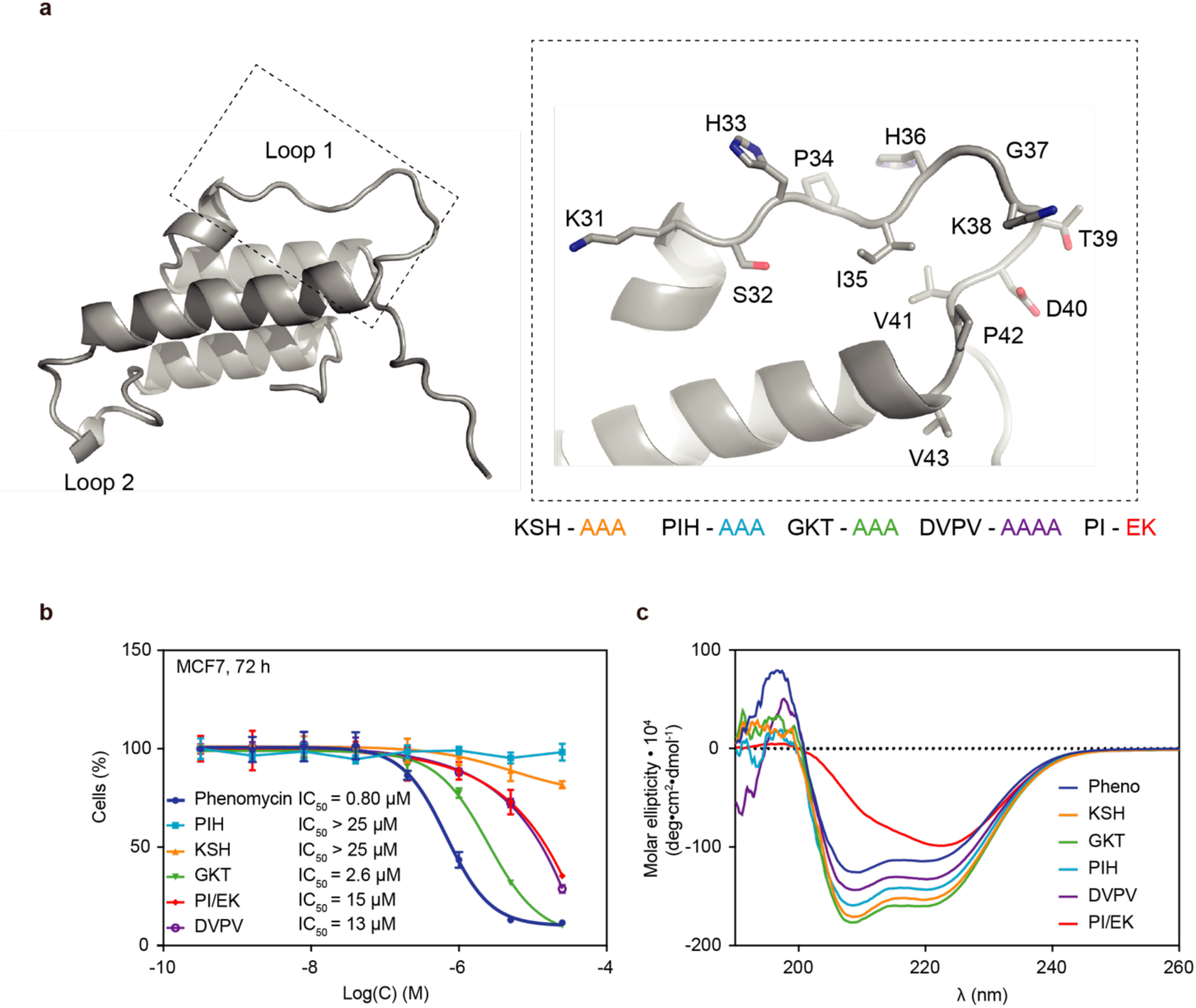
Assessing the role of loop 1 in cytotoxicity of phenomycin. (a) Zoom-in on the structure and sequence of loop 1. We generated separate mutants of triplets or quartets of alanine, for example the three amino acids KSH is substituted with AAA and results are shown in orange. (b) Inhibition of MCF7 cell viability, only the mutants where PIH or KSH is substituted with AAA lose all measurable toxicity. Data points are mean ± s.d. (*N*=3). Curves are representative from an experiment performed three times with consistent outcome. (c) CD measurements of phenomycin and mutants at 25°C. See also Figure S6.

## Discussion

The ability of phenomycin to inhibit translation was documented decades ago. Specifically, the initiation steps appeared to be affected at low concentrations of the protein whereas at higher concentrations elongation steps were also inhibited.^17^ It should be noted that these experiments were all conducted in cell free systems and the rapid (minutes) and potent (low nanomolar) inhibitory activities observed under such conditions stand in rather dramatic contrast to the activity of the protein in living cells, where toxicity typically manifests only over the course of several days at much higher concentrations. Indeed, whether the toxicity of phenomycin towards mammalian cells is truly mediated by a direct inhibition of protein synthesis cannot be conclusively determined based on the previous data. In this paper, we have provided several pieces of data that do support such a conclusion: (I) Using a fluorophore-conjugated phenomycin-derivative, we demonstrate that the protein accumulates in late endosomes via an uptake mechanism that appears in accord with endocytosis. We also found that a protein transfection agent could enhance toxicity of phenomycin. (II) We combined direct measurements of cellular protein synthesis using a puromycin-based assay with morphological profiling to show that phenomycin both blocked protein synthesis and mechanistically clustered closely to a panel of known small molecule translation inhibitors (cycloheximide, homoharringtonine, emetine) that directly bind and inhibit the ribosome. (III) In vitro binding assays based on BLI determined the affinity of biotin-conjugated phenomycin to the 80S human ribosome to be in the picomolar regime. Taken together, this points to a mechanism of action involving endosomal escape as the rate-limiting step and that even very low concentrations of the toxin in the cytosol can potently inhibit protein synthesis. This scenario would explain why we do not observe any significant cytosolic localization of the fluorophore-labelled phenomycin although we cannot formally rule out that a proteolytic event, that cleaves the N-terminal and thereby the fluorophore, precedes endosomal escape. We have generated the first structural insight into this class of bacterial proteins and by conducting systematic mutations we have pinpointed loop 1, in particular the region featuring a lysine (K31) and two surface exposed histidine residues (H33 and H36), as a critical component for activity. Histidines are well known to stack with nucleobases and can change their charge state around physiological pH. We suspect that this segment could be involved in the interaction with the ribosome, however, the very similar sequence present in carotomycin, which lacks growth-inhibitory and ribosome-binding activity, and our loop1/2-hybrid variants suggests that this segment alone is not sufficient. Either the specific spatial presentation or another structural element is also required for the potency of phenomycin. Our demonstration of the importance of loop 1 can also potentially explain an early observation that a derivative of phenomycin with one histidine residue carboxymethylated lost *in vivo* activity.^37^ This derivative was generated by exposure of phenomycin to iodoacetic acid followed by purification and amino acid analysis, however the modified histidine residue was not identified. Loop 1 contains two of the four histidines present in the protein and the other two remaining histidines are located in helix 3 towards the C-terminal. Despite several attempts we have thus far unfortunately not been able to determine the structure of a phenomycin-ribosome complex although further investigations towards this end are ongoing. Phenomycin, and its closests homologs, are examples of the emerging class of folded bacterial mini proteins^38^ that can facilitate potent and specific biological activities. While the ecological role of phenomycin seems apparent in terms of acting as a toxin towards e.g. fungi in the microenvironment, the functions of non-cytotoxic phenomycin homologs, such as carotomycin, still remains unknown and should be addressed in future investigations. In addition, the possibility for grafting^39^ bioactive peptides into the loops of phenomycin, to facilitate their intracellular delivery is also an interesting opportunity.

## Significance

Small proteins with stable folds that contain only a few secondary structure elements are referred to mini-proteins. Despite of their modest size — 50-100 amino acids or less — such proteins can mediate diverse biological functions and they constitute a highly active area of research. The discovery of new small proteins has been accelerated by genomic and proteomic technologies, but some were discovered using traditional natural product isolation procedures. The toxin phenomycin, the subject of this study, is one such example being initially isolated from a streptomyces strain more than 50 years ago and subsequently found to be an 89-residue protein. Phenomycin inhibits ribosomal protein synthesis and possesses antitumor activity in animal models. In our studies, we have found that phenomycin-like proteins are wide-spread in actinobacteria but that homologs, that do not share the toxicity of phenomycin, are also found in other bacterial phyla. We have solved the first structure of a member of this protein family which e.g. allowed the pinpointing of a short segment in one of the two loops that mediate the toxicity of phenomycin in mammalian cells. This insight may provide an actionable point-of-entry for future detailed mechanistic investigations on how phenomycin selectively perturbs the function of the eukaryotic ribosome. Our data also demonstrate that phenomycin must possess the ability to access the cellular cytoplasm through endosomes in order to reach its target. As endosomal escape is a central challenge for development of intracellular biological drugs, the understanding of how this naturally occurring protein achieves this key function is of strong interest.

## Acknowledgements

Katrine Hansen is acknowledged for technical assistance. This work was supported by the Carlsbergfoundation (grant CF17-0800 to T.B.P. and grant 2013_01_0566 to L.N.N), Independent research fund Denmark (Sapere aude 2 grant 6110-00600A to T.B.P.). Access to the 950 MHz NMR spectrometer at the Danish Center for Ultrahigh-Field NMR Spectroscopy (Ministry of Higher Education and Science grant AU-2010-612-181) is acknowledged.

## Author contributions

Conceptualization, T.B.P. and T.T.; Investigation, B.K.H, C.K.L, J.T.N., E.B.S., K.M.J., M.B., T.T. and L.B.V.; Writing – Original Draft, T.B.P. and T.T.; Writing – Review & Editing, T.B.P. and T.T.; Visualization: T.B.P., T.T., B.K.H., C.K.L., J.T.N., and E.B.S; Resources, R.K.F. and L.B.J.; Supervision, T.B.P., T.T., F.A.A.M., D.E.B., L.N.N. Funding Acquisition, T.B.P.

## STAR methods

### CONTACT FOR REAGENT AND RESOURCE SHARING

Further information and requests for resources and reagents should be directed to and will be fulfilled by the Lead Contact, Thomas B. Poulsen.

There are restrictions to the availability of the conjugated derivatives of phenomycin, carotomycin, and aquimarinamycin due to the lack of an external centralized repository for its distribution and our need to maintain the stock. We are glad to share the plasmids encoding the M3C mutants that enable conjugation with IA-alkyne.

### EXPERIMENTAL MODEL AND SUBJECT DETAILS

#### Cell culture

The human breast cancer cell line MCF7 (RRID: CVCL_0031, taken from 69-year-old female) was cultured in DMEM (Gibco, Cat#: 21969-035) supplemented with Ala-GLn (200 μM, Sigma-Aldrich, Cat#: G8541), Fetal Bovine Serum (FBS) (10%, Gibco, Cat#: A31609-02), Penicillin/Streptomycin (1%, Gibco, Cat#: 15140122) and insulin solution human (0.01 mg/ml, Sigma-Aldrich, Cat#: I9278). PANC-1 cells (RRID: CVCL_0480, taken from 56-year-old male) were cultured in DMEM supplemented with FBS (10%) Penicillin/Streptomycin (1%) and Ala-GLn (200 μM). BxPC3 cells (RRID: CVCL_0186, taken from a 61-year-old female) were cultured in RPMI 1640 (Gibco, Cat#: 61870-010) supplemented with FBS (10%) and Penicillin/Streptomycin (1%). Human lung carcinoma epithelial cell line HCC827 (RRID: CVCL_2063, taken from a 39-year-old female) was cultured in RPMI 1640 supplemented with FBS (10%) and Penicillin/Streptomycin (1%). Breast cancer cell line MDA-MB-231 (RRID: CVCL_0062, taken from 51-year-old female) was cultured in DMEM/F12 (Gibco, Cat#: 31331-028) supplemented with insulin (5 μg/ml), FBS (5%) and Penicillin/Streptomycin (1%). A549 cells (RRID: CVCL_0023, taken from 58-year-old male) cultured in Ham’s F-12K (Kaighn’s) (Gibco, Cat#: 21127-022) supplemented with FBS (10%) and Penicillin/Streptomycin (1%). The cervical tumor cell line SiHa (RRID: CVCL_0032, taken from 55-year-old female) was cultured as monolayers in MEM (Gibco, Cat#: 41090-028) supplemented with FBS (10%, ThermoFisher Scientific, Cat#: 12350273), Penicillin/Streptomycin (1%), 1% non-essential amino acids (Gibco, Cat#: 11140-035) and 1% sodium-pyruvate (Gibco, Cat#: 11360-039). HaCat cells (RRID: CVCL_0038, taken from 62-year-old male) were cultured in DMEM supplemented with FBS (10%) and Penicillin/Streptomycin (0.5%). Normal mouse hepatocyte cell line AML12 (RRID: CVCL_0140, taken from male) was cultured in DMEM/F12 medium supplemented with insulin (0.01 mg/ml), transferrin (5 μg/ml), selenium (5 ng/ml), dexamethasone (40 ng/ml), FBS (10%) and Penicillin/Streptomycin (1%). Human osteosarcoma cell line U2OS (RRID: CVCL_0042, taken from a 15-year-old female) was cultured in McCoy’s 5A (Sigma, Cat#: M9309) supplemented with FBS (10%, Gibco, Cat#: A31609-02) and Penicillin/Streptomycin (1%, Gibco, Cat#: 15140122).

Cells were grown in T75 flasks (Biolite, Thermo Scientific, Cat#: 130190) in a humidified atmosphere with 5% CO_2_ at 37 °C. Cells were passaged approximately every third day by trypsinization (Sigma-Aldrich, Cat#: T4049) followed by transfer of one-fifth of cells to a new flask. All cell cultures were handled aseptically in a laminar air flow bench and grown in sterile-filtered (Jetbiofil, Cat#: FPE204500) medium according to ATCC’s recommendations.

## METHOD DETAILS

### Cloning

All primers and some of the constructs were synthesized by Integrated DNA Technologies (IDT).

Dry intact genes codon optimized for *E. coli* from IDT (Table S4) were dissolved in ddH_2_O (10 ng/μL) and 200 ng were cleaved with NcoI (#FD0573) and XhoI (#FD0694) and purified (Thermo Scientific PCR purification kit #K0701). The vector (pETM-11) was cleaved with the same two restriction enzymes and purified using gel electrophoreses. Upon ligation with T4 DNA ligase (#EL0011), vector and insert were mixed in a 1:3 molar ratio using 50 ng of pETM-11 and incubated for 30 minutes at room temperature, before transforming chemically competent *E. coli* DH5α with ligation mixture. 3-5μl of the ligation mixture was transferred to DH5α (50μl) and incubated 30 min on ice, followed by heat shock (42°C) for 50 seconds and 2 min incubation on ice. SOC media (400μl) was used for rescue together with incubation (37°C, 45 min). Transformed cells were selected for by plating on LB agar containing kanamycin (50 μg/ml).

### Site-directed mutagenesis

Cysteine to alanine mutations were made by PCR with primers from IDT. Template DNA (pETM-11 with insert, 100 ng) was mixed in FD Buffer, dNTPs (0.2 mM), forward primer (0.5 μM), reverse primer (0.5 μM), DMSO (3%), and Phusion polymerase (0.02U/μL) (#F530S). The final PCR reaction was treated with DpnI (#FD1704), according to the manufactures protocol, for removal of original template.

### Protein expression

For protein expression *E. coli* BL21 (DE3) or LOBSTER^40^ chemically competent cells were transformed with pETM11 with insert and selected using kanamycin (50 μg/ml). Starter cultures were made from a single colony and used for inoculation of LB media (2 L) with kanamycin (50 μg/ml). After reaching an OD_600_ of ∼0.6, cells were induced with IPTG (0.5 mM) overnight at 18 °C. The next morning cells were harvested by centrifugation at 6000 rpm for 20 minutes at 4 °C. If the purification was not continued immediately after harvest, cell pellets were stored at -20 °C for up to one month. For the expression of labeled phenomycin for NMR studies, the *E. coli* BL21 (DE3) was grown in minimal medium (3 × 1 L) containing ^15^NH_4_Cl (1 g/L) and U-^13^C glucose (3 g/L). The culture was incubated at 37°C and allowed to reach an OD_600_= 0.75 before cooling on ice, inducing with IPTG, and incubation overnight at 18°C. See further details below.

### Protein purification

Proteins were purified according to their His_6_-tag using a His Excel column from GE Healthcare attached to a Äkta Start system. Cell pellets were suspended in buffer A (50 mM HEPES, 500 mM NaCl, 10 mM Imidazole, pH 7.5) (2 mM β-mercaptoethanol was added to buffer A, when the cysteine mutants were purified), sonicated 3×5 minutes (4 cycles, ∼ 60% power) and incubated on ice between the runs to prevent sample heating. The lysate was clarified by centrifugation at 15,000 rpm for 40 minutes at 4°C. The supernatant was filtered (0.22 μm) before loading on a pre-equilibrated column and washed with 8-10 bed volumes of buffer A. Protein was then eluted using a gradient of buffer B (50 mM HEPES, 500 mM NaCl, 400 mM Imidazole, pH 7.5) (2 mM β-mercaptoethanol was added to buffer B, when cysteine mutants were purified) to 150 mM and remaining protein was stepped out with 100% B. The eluted protein was dialyzed against 50 mM HEPES, 500 mM NaCl, and 10 mM Imidazole, pH 7.5 and cleaved with TEV protease in a 1:10 w/w ratio for His_6_-tag removal. The mixture was applied to a His Excel column to remove the cleaved His6-tag, the TEV-protease, and unspecific binders. The purified protein was collected in the flow through. Eluted protein was dialyzed against 50 mM HEPES and 200 mM NaCl pH 7.5 for removal of imidazole and subsequently concentrated using Amicon^®^ Ultra-4 spin columns (# UFC800308). The final protein concentration was calculated from the absorbance at 280 nm (NanoDrop) with the predicted extinction coefficient calculated in CLC Main Workbench. The purity was assessed by SDS-PAGE.

### Cell viability assays

MCF7 cells were seeded in black 96-well plates (Thermo Scientific, Cat#: 137103) at a density of 2000 cells/well in complete growth medium (75 μL/well) and allowed to adhere overnight. The next day, each protein were five-fold serial diluted in PBS as 200X concentrations and further diluted in media to 4X solutions (8 dilutions). The proteins were given to cells by the addition of 25 μL/well in triplicates resulting in 1X solutions (final concentrations of 25 μM-0.32 nM) in 100 μL/well. The cells were then incubated at 37 °C in a humidified atmosphere containing 5% CO_2_. 90 minutes before end of treatment, all cells were given CellTiter Blue (20 μL/well, Promega, Cat#: G8081) and incubated for the remaining time of the experiment. The viability of the cells was assessed by measuring fluorescence (552 ± 10 nm excitation; 598 ± 10 nm emission) in a Tecan Spark 10 M multimode plate reader. Data was corrected for background fluorescence, normalized to the average of the two lowest concentrations of the proteins and fitted to a four-parameter nonlinear regression using GraphPad Prism version 7.0 for Mac OS X, GraphPad Software, La Jolla California USA.

For fluorophore-labelled phenomycins, cell viability was analyzed by light microscopy. MCF7 cells were seeded in a clear 96-well plate (Thermo Scientific, Cat#: 167008) as described above and treated with fluorophore-labelled phenomycins (1 and 5 μM) for 72 hours at 37 °C in a humidified atmosphere containing 5% CO_2_. After treatment, optical images of the cells were taken by light microscopy.

### Alkylation with an iodoacetamide-derivative having a terminal alkyne (IA-alkyne)

Cysteine mutants were conjugated with an iodoacetamide-derivative for subsequent click chemistry. Protein (15 mg) was reduced with TCEP (5 mM) for 10 min and mixed with IA-alkyne (70 μl of 200 mM) and MeCN (300 μl), the reaction was incubated overnight at room temperature. Protein tagged with IA-alkyne was purified by reverse phase HPLC on a semipreparative Vydac 5 μm C4 300 Å protein column (10×250 mm, 7 ml/min) with a mobile phase of H_2_O + 0.1% TFA (buffer A) and MeCN + 0.1 % TFA (buffer B). Samples were loaded on a pre-equilibrated (20% A) column and proteins were eluted using a gradient elution profile as follows: 20-40% B, 30 min – 40% B, 4 min – 40-100% B, 1 min – 100% B, 4 min, and absorbance was monitored at 210 nm. Fractions were collected and analyzed on an autoflex speed MALDI-TOF mass spectrometer (Bruker Daltonics) for a positive alkylation using a sinapinic acid (SA) matrix. Fractions with alkyne labelled proteins were combined, lyophilized and stored in the freezer.

### Copper(I)-catalyzed click reaction

Copper-catalyzed azide-alkyne cycloaddition (CuAAC) was performed to conjugate fluorophores or biotin to the proteins. The alkyne labelled proteins were dissolved in 0.1 M phosphate buffer with 100 mM NaCl and 5 mM aminoguanidine (pH 7.2) to 2 mg/ml (approximately 250 μM alkyne). The proteins were mixed with reagents for CuAAC: Final concentrations: CuSO_4_ (1.25 mM), Tris(3-hydroxypropyltriazolylmethyl)-amine (THPTA) (5.9 mM, Sigma-Aldrich, Cat#: 762342) and sulfo-Cy5-azide (Lumiprobe, Cat#: A3330), FAM-azide (Lumiprobe, Cat#: C4130) or Azide-PEG3-biotin (Sigma-Aldrich, Cat#: 762024) (2 eq., 500 μM). The solutions were vortexed and the click reactions initiated by addition of (+)-sodium L-ascorbate (4.2 mM, Sigma-Aldrich, Cat#: A7631). The reactions incubated for 6 hours at room temperature and subsequently purified by reverse phase HPLC as described above. Fractions were analyzed by MALDI-TOF and fractions with eluted protein were combined and lyophilized. The proteins were afterwards dissolved in 0.1 M phosphate buffer with 100 mM NaCl.

### Cellular uptake studies of fluorophore-labelled proteins

Nunc Lab-Tek 8-well chambered coverglass (Thermo scientific, Cat#: 155411) was washed twice in PBS and coated with poly-D-lysine hydrobromide (50 μg/mL, Sigma-Aldrich, Cat#: P7405) for 2 hours at room temperature. Afterwards, the slide was washed twice with PBS, and MCF7 cells were seeded at a density of 12,500 cells/well in complete medium (200 μL/well) and the slide incubated at 37 °C in a humidified atmosphere containing 5% CO_2_. The next day, fluorophore-labelled proteins were diluted in complete growth media and the solutions filtered to remove aggregates. The cell culture media were removed and replaced with the growth media containing the proteins (150 μL/well). The cells were incubated at 37 °C in a humidified atmosphere containing 5% CO_2_ for 12 hours. 15 minutes before end of treatment, Hoechst 33342 (12.3 mg/mL, Thermo Scientific, Cat#: 62249) was diluted to a 4X solution in media and added to all wells (1 μg/mL, 50 μL/well) for staining the nuclei of the cells. After treatment, the cells were washed three times with growth medium (200 μL) and new growth medium (200 μL) was added before live imaging. Images were acquired using a Zeiss LSM700 confocal laser scanning microscope (CLSM) equipped with a 63x NA1.4 Plan-Apochromat oil immersion objective (Zeiss) and excitation with lasers 488 nm and 639 nm.

### GFP-Rab7 co-localization studies

Cells were grown on poly-D-lysine coated 8-well chambered coverglasses at a density of 12,500 cells/well and allowed to adhere overnight. The cells were transfected with GFP-Rab7 plasmids (Addgene plasmid #12605) using a Lipofectamine 3000 Reagent kit (Thermo Scientific, Cat#: L3000001) according to the manufacturers’ protocol. Three days after transfection, the cell media were removed in all wells and replaced with filtered sulfo-Cy5-labelled phenomycin (5 μM) in complete growth medium (150 μL/well). After 12 hours treatment, Hoechst 33342 (1 μg/mL) was added for 15 min. The cells were then washed twice with media (200 μL/well) and new growth media (200 μL/well) was added before live imaging with CLSM.

### Inhibition of cellular uptake by 4°C incubation

MCF7 cells were seeded in four wells of two 8-well chambered coverglasses (15,000 cells/well, 150 μL/well) in complete growth medium and allowed to adhere overnight. The next day, one of the coverglasses were placed in the fridge (4°C, 30 min) prior to treatment. Transferrin conjugated with Alexa Fluor 555 (25 μg/ml, ThermoFisher Scientific, Cat#: T35352) or FAM-phenomycin (20 μM) were added in duplicates to wells of each of the two coverglasses (200 μL/well) and one coverglass were placed at 4 °C and the other at 37 °C for 4 hours. 15 min before end of treatment, Hoechst 33342 were added as 5x solutions (50 μL/well, final concentration: 1 μg/mL). After incubation, the cells were washed three times in cold or warm medium and new medium were added before CLSM imaging.

### BioPORTER protein delivery

Intracellular delivery of phenomycin into living cells was performed using the BioPORTER Protein Delivery Reagent QuickEase Kit (AMS Biotechnology (Europe), Cat#: BP502424). MCF7 cells were seeded in a black 96-well plate (2000 cells/well, 75 μL/well) in complete growth medium and allowed to adhere overnight. The next day, phenomycin (1.05 mM) was diluted to 25x solutions in PBS (final concentrations of 200 nM, 5 μM and 25μM). 40 μL of the diluted protein solutions or PBS were either transferred to new eppendorf tubes or used to hydrate QuickEase tube of BioPORTER Reagent. The BioPORTER samples were gently pipetted up and down five times, incubated for 5 min at room temperature and then gently vortexed (3-5 sec). 460 μL serum-free growth medium was added to all tubes to get a final volumen of 500 μL. The culture medium was removed from the cells, and the cells washed once with serum-free growth medium before addition of 50 μL serum-free growth medium to each well. The BioPORTER:protein mix or protein samples were added to cells in triplicates and the cells incubated at 37 °C in a humidified atmosphere containing 5% CO_2_. After 4.5 hour incubation, 100% FBS (11 μL/well) was added to each well and the cells were reincubated for the remaining time of the experiment (72 hours). The rest of the experiment was carried out as a regular cell viability assay.

### Ribosomal purification

Bacterial ribosomes were purified from 20 g *Thermus thermophilus* cell pellet (BFF, Athens Georgia, USA) with all steps carried out at 4ºC. Cells resuspended in buffer A (20 mM HEPES/KOH pH 7.5, 100 mM NH_4_Cl, 10 mM Mg(OAc)_2_, 0.5 mM EDTA, 1 mM DTT and 0.1 mM benzamidine) were lysed by three rounds of high-pressure homogenisation with a backpressure of 20,000 psi. Lysate was treated with RNase-free DNaseI (1U/μL, ThermoFisher, #EN0521) before clearing by centrifugation for 45 min at 30,000 x g. Crude ribosomes were pelleted through a sucrose cushion (20 mM HEPES/KOH pH 7.5, 500 mM KCl, 10 mM Mg(OAc)_2_, 0.5 mM EDTA, 37% w/v sucrose, 1 mM DTT) by ultracentrifugation for 17 hours at 125,171 x g. Pellets were gently washed in buffer CT (20 mM HEPES/KOH, 400 mM KCl, 10 mM Mg(OAc)_2_, 1.5 M (NH_4_)_2_SO_4_, 1 mM DTT) before resuspended in buffer CT (5 mL) and loaded onto a butyl-ToyoPearl column (TOSOH) equilibrated in buffer C. Column was washed for 1 column volume with 50% buffer CT and 50% buffer D (same as buffer CT without (NH_4_)_2_SO_4_) and ribosomes eluted with a 20 column volume linear gradient into buffer D. Eluted ribosomes were pelleted by ultracentrifugation for 17 hours at 125,171 x g. Ribosomal pellets were resuspended in buffer RE (10 mM HEPES/KOH pH 7.5, 50 mM KCl, 10 mM NH_4_Cl, 10 mM Mg(OAc)_2_, 0.25 mM EDTA, 1 mM DTT) and placed onto linear sucrose gradient (5-20% w/v) prepared in buffer RE. Ribosomes were sedimented by ultracentrifugation for 17 hours at 31,383 x g. Gradients were fractionated bottom-to-top and ribosome-containing fractions were pooled and ultracentrifuged for 17 hours at 125,171 x g to pellet ribosomes. The final ribosome pellets were dissolved in buffer G (5 mM HEPES/KOH pH 7.5, 50 mM KCl, 10 mM NH_4_Cl, 10 mM Mg(OAc)_2_, 1 mM DTT) to a final concentration of 20 mg/mL. Aliquots were snap-frozen in liquid nitrogen and stored at -80ºC.

Yeast ribosomes were purified from *Saccharomyces cerevisiae* cells not expressing Stm1 protein (strain JD1559 from J. Dinman’s lab at University of Maryland).^41^ All steps were carried out at 4ºC. Approximately 5 g of cells were resuspended in buffer M (10 mL) (30 mM HEPES/KOH pH 7.5, 50 mM KCl, 10 mM MgCl_2_, 8.5% w/v mannitol, 0.5 mM EDTA, 2 mM DTT, protease inhibitor cocktail (Sigma)) and RNasin (100μl, Promega) added. The resuspended cells were lysed in a SS34 centrifugation tube by vortex with of glass beads (13 g) (400-600 micron) for one minute and two minutes pause. This was repeated five times. Crude lysate was obtained by centrifugation for 2 min at 20,000 x g and the supernatant further cleared by centrifugation for 10 min at 30,000 x g. PEG 20,000 (Hampton Research) was added to the clear supernatant to a final concentration of 4.5% w/v and the solution was left on ice for 5 min before centrifuged for 5 min at 20,000 x g. Supernatant was transferred to a new centrifugation tube and KCl concentration adjusted to 130 mM and incubated for 5 min on ice. PEG 20,000 was further added to a final concentration of 8.5% w/v and the solution left on ice for 10 min. Precipitated ribosomes were pelleted by centrifugation for 10 min at 20,000 x g. The resulting pellet was resuspended to 7 mg/mL in buffer M2 (buffer M made with 150 mM KCl). The ribosome suspension was carefully place onto linear sucrose gradient (10-30% w/v) prepared in buffer A (20 mM HEPES/KOH pH 7.5, 120 mM KCl, 8.3 mM MgCl_2_, 0.3 mM EDTA, 2 mM DTT). Ribosomes were sedimented by ultracentrifugation for 15 hours at 37,348 x g. The gradients were fractionated bottom-to-top and peak fractions of ribosome pooled and dialyzed against buffer C (20 mM HEPES/KOH pH 7.5, 50 mM KCl, 5 mM MgCl_2_, 10 mM NH_4_Cl, 1 mM DTT) overnight. The dialyzed solution was loaded onto a column packed with 24 mL cysteine-linked Sulfolink resin (Pierce)^42^ equilibrated in buffer C. The column was washed in 1 column volume buffer C and bound ribosomes eluted with a 10 column-volume linear gradient into buffer C800 (buffer C made with 800 mM KCl). Two peaks eluted from the column around 280 mM KCl and 380 mM KCl. Fractions of the respective peaks were pooled separately. Both ribosomes solutions were buffer exchanged into buffer C using concentrator spin columns with a 100,000 cut-off. SDS-PAGE analysis suggested ribosomes of the first peak to be eEF2 bound, while ribosomes of the second elution peak were “empty” ribosomes. Only the empty ribosomes were used further on. Final concentration of yeast ribosomes was 10 mg/mL. Aliquots were snap-frozen and stored at -80ºC.

Human ribosomes were purified from HEK-293F cell pellet (40 g) (kindly provided by Prof. Gregers Rom Andersen, AU). All steps were carried out on ice or a 4ºC. Cells were lysed in buffer L (20 mM HEPES/KOH pH 7.2, 50 mM KCl, 5 mM MgCl_2_, 10% v/v glycerol, 0.5% v/v NP-40, 1 mM DTT, protease inhibitor cocktail pill (Sigma)) for 40 min. Lysate was cleared by centrifugation for 30 min at 30,000 x g. Crude ribosomes were pelleted through a sucrose cushion (50 mM HEPES/KOH pH 7.5, 150 mM KCl, 5 mM MgCl_2_, 10 mM NH_4_Cl, 60% w/v sucrose, 1 mM DTT) by ultracentrifugation for 3 hours at 125,171 x g. The resulting pellet was resuspended in buffer B (50 mM HEPES/KOH pH 7.5, 100 mM KCl, 3 mM MgCl_2_, 1 mM DTT) and the solution placed onto a linear sucrose gradient (15-40% w/v) prepared in buffer B. Ribosomes were sedimented by ultracentrifugation for 16 hours at 42,874 x g. Gradients were fractionated bottom-to-top and ribosome fractions were collected, pooled and dialyzed overnight against 1 L buffer C (20 mM HEPES/KOH pH 7.5, 50 mM KCl, 5 mM MgCl_2_, 10 mM NH_4_Cl, 1 mM DTT). The dialyzed ribosome solution was loaded onto a column packed with 24 mL cysteine-linked Sulfolink resin (Pierce)^42^ equilibrated in buffer C. The column was washed in 1 column volume buffer C and bound ribosomes eluted with a 10 column-volume linear gradient into buffer C800 (buffer C made with 800 mM KCl). Elution fractions containing ribosomes were pooled and buffer exchanged into buffer C on a concentrator spin column with a 100,000 cut-off. Final concentration of human ribosome was 5.9 mg/mL. Aliquots were snap-frozen in liquid nitrogen and stored at -80ºC.

Plant ribosomes were purified from commercially available wheat germ. All steps were carried out on ice or at 4ºC. 100 gr of wheat germ were frozen in liquid nitrogen and ground into a fine powder using a mortar and pestle. Powdered wheat germ was dissolved in ice-cold plant extraction buffer (1:2 v/v) (50 mM HEPES/KOH pH 7.5, 400 mM KCl, 5 mM MgCl_2_, 17% w/v sucrose, 1 mM DTT) and further homogenized in a blender. The suspension was cleared by centrifugation for 30 min at 30,000 x g. The supernatant was added 0.1 x volume of Triton X-100 (20% v/v) and centrifuged again for 20 min at 30,000 x g. Crude ribosomes of the supernatant were pelleted through a sucrose cushion (50 mM HEPES/KOH pH 7.5, 150 mM KCl, 5 mM MgCl_2_, 10 mM NH_4_Cl, 60% w/v sucrose, 1 mM DTT) by ultracentrifugation for 3 hours at 125,171 x g. The pellet was dissolved in resuspension buffer (50 mM HEPES/KOH pH 7.5, 100 mM KCl, 3 mM MgCl_2_, 1 mM DTT) and placed onto a linear sucrose gradient (15-40%). Ribosomes were sedimented by ultracentrifugation for 16 hours at 42,874 x g. Gradients were fractionated bottom-to-top and ribosome fractions collected, pooled and dialyzed against 1 L buffer C overnight. Dialyzed ribosomes were further purified with chromatographic step described above for human ribosomes. Eluted ribosomes were pooled and buffer exchanged into buffer C using a concentrator spin column with a 100,000 cut-off. Final concentration of plant ribosomes was 6 mg/mL. Aliquots were snap-frozen in liquid nitrogen and stored at -80ºC.

### Biolayer Inferometry (BLI)

BLI experiments were performed on Octet Red96 (FortéBio) at 30°C and shaking at 1000 rpm using black non-transparent 96-well plates. The sensors were equilibrated in kinetic buffer (50 mM HEPES, 0.1 % BSA (Sigma, #A3912), 5 mM MgCl_2_, 0.05 % TWEEN in PBS for 10 minutes. Biotinylated protein dissolved in kinetic buffer at 0.5 μg/ml was loaded on Streptavidin BLI sensors (SA, FortéBio, 18-5019) for 5 minutes and then washed for 10 minutes in kinetic buffer. Associations of human ribosomes (4-0.5 nM), bacterial ribosomes (4-0.25 nM), yeast ribosomes (8-0.25 nM) and plant ribosomes (4-1 nM) diluted in kinetic buffer were monitored for 10-15 minutes followed by 5-10 minutes of dissociation. The kinetic constants were determined with FortéBio Data Analysis software using a 1:1 Langmuir binding model.

### Assay definition

Dip and Read Streptavidin sensors (FortéBio) were hydrated in kinetic buffer 10 minutes before use. A baseline was established (600 sec) before loading with biotinylated protein (300 sec), baseline (600 sec), then association with ribosome (900 sec) and dissociation (1800/3600 sec). Data was analyzed using FortéBio Data Analysis software.

### Circular Dichroism (CD) spectrometry

The buffer was changed to CD buffer (50 mM NaPO_4_ (pH 7.4), 20 mM NaCl) using 2 ml Zeba™ Spin Desalting Columns, 7 kDa cut-off (Thermo Scientific, #89890). The protein concentration was adjusted to 0.5 mg/ml in CD buffer (250 μL). CD spectra were recorded on a JASCO Corp. J-810 with at 25°C with a bandwidth of 2 nm, a scanning speed of 100 nm/min and an accumulation of three scans. For the UV CD, a scan range of 190-260 nm and a cuvette with a path-length of 0.1 mm was used. The background from the buffer was subtracted, and data was normalized to their molecular weight in order to compare the spectra. CD data were recorded in mdeg and converted to molar ellipticity (deg*cm^2^*dmol^-1^). To test the stability the proteins were subsequently scanned with increasing temperature (25-95°C), one degree per minute with a fixed wavelength at 222 nm.

### Morphological profiling

4000 U2OS cells were seeded into the inner 60 wells of a 96-well plate with optical bottom (Corning cat. no. 3603) in 50 μL full growth medium and incubated (37 °C, 5% CO_2_, humid) for 24 h. Peptide compounds were diluted to 30X in PBS followed by dilution to 4X in media before addition of 25 μL compound to each well followed by addition of 25 μL 2% DMSO (final DMSO = 0.5%). Translation inhibitors were diluted to 200X in media followed by dilution to 4X before addition of 25 μL compound to each well followed by addition of 25 μL 13% PBS. All treatments are performed in quadruplicate with 12 DMSO negative controls per replicate. After 24 h 75 μL medium was removed and replaced with 75 μL medium containing 500 nM MitoTracker Deep Red (final C = 325 nM) and plates were incubated in the dark for 30 min. Wells were then aspirated and 75μL medium were added, before adding 25 μL 16 % paraformaldehyde (Electron Microscopy Sciences 15710-S) (final PFA = 4 %) and incubating in the dark for 20 min. Plates were washed once with 1X HBSS (Invitrogen 14065-056) and 75 μL 0.1 % (vol/vol) Triton X-100 (BDH 306324N) in 1X HBSS was added and incubated for 15 min in the dark. Plates were washed twice with 1X HBSS before addition of 75 μL multiplex staining solution (final concentrations: 1.5 μg/mL WGA-AF555; 35 μg/mL Concanavalin-AF488; 5 μL/mL Phalloidin-AF568; 3 μM SYTO 14; 5 μg/mL Hoechst 33342 in 1 % (wt/vol) BSA (Sigma-Aldrich A9647) and incubated for 30 min in the dark. Plates were then washed three times with 1X HBSS, with no final aspiration and imaged immediately in a Zeiss Celldiscoverer 7 automated microscope. The imaging settings can be seen in Table S2. 9 sites are imaged in each well.

Images were analysed in Cell Profiler 2.1.1 using previously published pipelines^25^. First images were illumination corrected followed by extraction of quality control parameters before the final analysis. Single-cell profiles were exported to an MySQL database and the data was processed in R 3.6.1 using previously published scripts^25^ affording the final per-treatment profiles. Data processing consists of removal of incomplete data points and normalization using a Z-score.

Profiles were compared using Pearson correlations and clustered using hierarchical clustering with average linkage and the Pearson correlation as the distance metric.

### NMR experiments

Protein for NMR spectroscopy was generated as described above, except that during culturing minimal media was used containing 3 g/L U-^13^C-glucose and 1 g/L ^15^N-ammonium chloride. NMR samples contained 1mM U-(^13^C/^15^N)-labelled Phenomycin in 10 mM phosphate buffer pH 3.25 with 5% D_2_O. 2D ^1^H–^13^C HSQC spectra, recorded at pH 6, are basically superimposable, indicating no change in structure in this pH range.

Backbone resonance assignments were obtained using 3D triple resonance NMR spectroscopy on a 500 MHz spectrometer (11.7 T) with a shielded-gradient TXI probe (Bruker, Inc). The sequential assignment followed standard protocols to match ^13^Cα and ^13^Cβ chemical shifts of neighbouring amino acids from strips drawn from 3D HNCACB, using GAMES_ASSIGN.^43^ Simultaneously, the backbone ^13^C’ carbonyl chemical shifts were linked by use of 3D HNCO and 3D HN(CA)CO spectra (see below). Completion of the aliphatic side chain ^13^C chemical shift assignment was obtained by 3D (H)C-TOCSY-C(CO)NH, while the aliphatic ^1^H side chain chemical shifts were collected from 3D H(C)-TOCSY-C(CO)NH and 3D HBHA(CO)NH spectra. Finally, side chain ^1^H and ^13^C chemical shifts were connected by inspection of 2D ^1^H–^13^C HSQC and 3D H(C)CH spectra.

Subsequently, 3D ^13^C-^1^H-HSQC-NOESY and ^15^N-^1^H-HSQC-NOESY spectra were recorded at 950 MHz. ^13^C-^1^H-HSQC-NOESY was first employed to establish the assignment of aromatic side chain ^1^H and ^13^C chemical shifts by connecting the aliphatic and aromatic assignments via the strong proton-protein NOEs between β and δ protons. The backbone assignment was complete. Side chain assignment coverage was also complete; All non-labile hydrogens were assigned, as were chain H–N in Asn, Gln, Trp and Arg (Nε–Hε).

### NMR structure determination and analysis of Phenomycin

The structure was calculated using distance constraints derived from 3D ^13^C-^1^H-HSQC-NOESY and ^15^N-^1^H-HSQC-NOESY spectra and dihedral angle constraints derived from the chemical shift using TALOS-N.^44^ The torsion angle predictions classified as “good” by TALOS-N were applied yielding 150 torsion angle restraints. The automatic simultaneous iterative structure refinement and NOE assignment within Cyana^45^ was applied in 32 parallel runs to produce 32 ensembles of structure and the 20 ensembles with lowest energy were selected. Consensus assigned NOE distance restraints were identified as restraints selected by Cyana in at least 5 of the 20 best ensemble runs. This procedure resulted in 1901 total validated distance restraints of which 543 were long range (residue number difference > 4). The final structure was recalculated using these validated 1901 distance restraints and choosing the 20 lowest energy structures from those 1000 calculated structures. Secondary structure elements were identified with Stride^46^ and the quality of the structure was evaluated using Cyana and the Protein Structure Validation Suite.^47^

## QUANTIFICATION AND STATISTICAL ANALYSIS

Where appropriate data is represented as mean ± s.d. Exact *N* values are stated for each experiment in the legends. *P*-values (Fig. 1c, Fig. 2d, Fig. 2e) were calculated in Graphpad Prism 8.2.0 (Students t-test, unpaired).

Hierarchical clustering was performed with the Pearson correlation as the distance metric and average linkage in R 3.6.0.

## DATA AND CODE AVAILABILITY

Per-treatment morphological profiles and the correlation matrix is uploaded on Mendeley Data (DOI: doi.org/10.17632/772b973hhz.1).

All scripts and code necessary for performing image analysis, feature extraction and data processing has been published previously (DOI: http://dx.doi.org/10.1016/j.bmc.2019.03.052). Images and single-cell profiles can be obtained upon request.

NMR chemical shift assignments for phenomycin can be obtained online from the Biological Magnetic Resonance Database (BMRB, http://www.bmrb.wisc.edu/) under accession number BMRB: 50088.

The accession number for the phenomycin structure is PDB: 6TKT.

## SUPPLEMENTARY INFORMATION

Figure S1. Supplementary viability and morphological profiling data.

Figure S2. Modified phenomycin maintains toxicity in mammalian cells and its cellular uptake is enhanced over time but can be impaired at low temperature.

Figure S3. Supplementary illustrations of the structure of phenomycin.

Figure S4. Protein alignment of phenomycin and enomycin with predicted proteins from other Streptomyces and Kitasatospora.

Figure S5. Stability and binding of phenomycin and homologs.

Figure S6. Detailed investigations of the two loop regions found in phenomycin.

Table S1. Overview of different proteins used in the study. Related to Figure 1-6 and Figure S1-S6. Table S2: Imaging settings for morphological profiling. Related to Figure 1 and Figure S6.

Table S3. Restraint and structure statistics for phenomycin. Related to Figure 3 and Figure S3. Table S4.

## References

1 Spradlin, J.N., Hu, X., Ward, C.C., Brittain, S.M., Jones, M.D., Ou, L., To, M., Proudfoot, A., Ornelas, E., Woldegiorgis, M. et al. (2019). Harnessing the anti-cancer natural product nimbolide for targeted protein degradation. Nat. Chem. Biol. 15, 747–755.

2 Velkov, T., Gallardo-Godoy, A., Swarbrick, J.D., Blaskovich, M.A.T., Elliott, A.G., Han, M., Thompson, P.E., Roberts, K.D., Huang, J.X., Becker, B. et al. (2018). Structure, function, and biosynthetic origin of octapeptin antibiotics active against extensively drug-resistant gram-negative bacteria. Cell Chem. Biol. 25, 380–391.

3 Filipuzzi, I., Steffen, J., Germain, M., Goepfert, L., Conti, M.A., Potting, C., Cerino, R., Pfeifer, M., Krastel, P., Hoepfner, D. et al. (2017). Stendomycin selectively inhibits TIM23-dependent mitochondrial protein import. Nat. Chem. Biol. 13, 1239–1244.

4 Kling, A. et al. (2015). Targeting DnaN for tuberculosis therapy using novel griselimycins. Science 348, 1106–1112

5 Jacobsen, K.M., Villadsen, N.L., Tørring, T., Nielsen, C.B., Salomón, T., Nielsen, M.M., Tsakos, M., Sibbersen, C., Scavenius, C., Nielsen, R. et al. (2018). APD-containing cyclolipodepsipeptides target mitochondrial function in hypoxic cancer cells. Cell Chem. Biol. 25, 1337–1349.

6 Villadsen, N.L., Jacobsen, K.M., Keiding, U.B., Weibel, E.T., Christiansen, B., Vosegaard, T., Bjerring, M., Jensen, F., Johannsen, M., Tørring, T., and Poulsen, T.B. (2017). Synthesis of ent-BE-43547A1 reveals a potent hypoxia-selective anticancer agent and uncovers the biosynthetic origin of the APD-CLD natural products. Nat. Chem. 9, 264–272.

7 Tsakos, M., Clement, L.L., Schaffert, E.S., Olsen, F.N., Rupiani, S., Djurhuus, R., Yu, W., Jacobsen, K.M., Villadsen, N.L., and Poulsen, T.B. (2016). Total synthesis and biological evaluation of rakicidin A and discovery of a simplified bioactive analogue. Angew. Chem. Int. Ed. 55, 1030–1035.

8 Nakamura, S., Yajima, T., Hamada, M., Nishimura, T., and Ishizuka, M. (1967). A new antitumor antibiotic, phenomycin. J. Antibiot. 20, 210–216.

9 Nishimura, T. (1968). Activity of phenomycin against transplantable animal tumors. J. Antibiot. 21, 106–109.

10 Muramatsu, R., Abe, S.-I., Hayashi, H., Yamaguchi, K., Jinda, K., Sakano, K.-I., Inouye, K., and Nakamura, S. (1991). Complete amino acid sequence of phenomycin, an antitumor polypeptide antibiotic. J. Antibiot. 44, 1222–1227.

11 Nishimura, T. (1968). Mechanism of action of phenomycin, a tumor-inhibitory polypeptide. J. Antibiot. 21, 110–118.

12 Suhara, Y., Ishizuka, M., Naganawa, H., Hori, M., Suzuki, M., Okami, Y., Takeuchi, T., and Umezawa, H. (1963). Studies on enomycin, a new antitumor substance. J. Antibiot. 16, 107–108.

13 Mizuno, S., Nitta, K., and Umezawa, H. (1967). Mode of Action of Enomycin, an Antitumor Antibiotic of High Molecular Weight. I. Inhibition of protein synthesis. J. Biochem. 61, 373–381.

14 Sakata, N., Oka, T., Ikeno, S., and Hori, M. (1994). Nucleotide sequence of the phenomycin gene from Streptoverticillium baldacci Ma564-C1. J. Antibiot. 47, 370–371.

15 Takeuchi, S., Oka, T., Sakata, N., S-Tuchiya, K, Hayashi, H., and Hori, M. (1997). Cloning of the enomycin structural gene from Streptomyces mauvecolor and production of recombinant enomycin in Escherichia coli. J. Antibiot. 50, 27–31.

16 Mizuno, S., Nitta, K., and Umezawa, H. (1967) Mode of Action of Enomycin, an Antitumor Antibiotic of High Molecular Weight. II. Preferential binding of enomycin to mammalian ribosomes. J. Biochem. 61, 382–387.

17 Yamaki, H., Nishimura, T., Kubota, K., Kinoshita, T., Tanaka, N. (1974) Selective inhibition of initiation of globin synthesis by phenomycin. Biochem. Biophys. Res. Commun. 59, 482–488.

18 Mizuno, S., Umezawa, H. (1976). Inhibition of the initial dipeptide synthesis of globin chains by the antibiotic enomycin in the reticulocyte lysate. J. Antibiot. 24, 309–315.

19 Bechara, C., and Sagan, S. (2013). Cell-penetrating peptides: 20 years later, where do we stand? FEBS Lett. 587, 1693–1702.

20 Bruce, V.J., McNaughton, B.R. (2017). Inside Job: Methods for Delivering Proteins to the Interior of Mammalian Cells. Cell Chem. Biol. 24, 924–934.

21 Walsh, M.J., Dodd, J.E., and Hautbergue, G.M. (2013). Ribosome-inactivating proteins. Virulence 4, 774–784.

22 Metelev, M., Osterman, I.A., Ghilarov, D., Khabibullina, N.F., Yakimov, A., Shabalin, K., Utkina, I., Travin, D.Y., Komarova, E.S., Serebryakova, M., et al. (2017). Klebsazolicin inhibits 70S ribosome by obstructing the peptide exit tunnel. Nat. Chem. Biol. 13, 1129–1136.

23 Mardirossian, M., Grzela, R., Giglione, C., Meinnel, T., Gennaro, R., Mergaert, P., and Scocchi, M. (2014). The host antimicrobial peptide Bac71-35 binds to bacterial ribosomal proteins and inhibits protein synthesis. Chem. Biol. 21, 1639–1647.

24 Bray, M.-A., Singh, S., Han, H., Davis, C.T., Borgeson, B., Hartland, C., Kost-Alimova, M., Gustafsdottir, S.M., Gibson, C.C., and Carpenter, A.E. (2016). Cell Painting, a high-content image-based assay for morphological profiling using multiplexed fluorescent dyes. Nat. Protoc. 11, 1757–1774.

25 Svenningsen, E. B., and Poulsen, T. B. (2019) Establishing cell painting in a smaller chemical biology lab - A report from the frontier. Bioorg. Med. Chem. 27, 2609–2615.

26 Carpenter, A.E., Jones, T.R., Lamprecht, M.R., Clarke, C., Kang, I.H., Friman, O., Guertin, D.A., Chang, J.H., Lindquist, R.A., Moffat, J., et al. (2006). CellProfiler: image analysis software for identifying and quantifying cell phenotypes. Genome Biol. 7, R100.

27 Schenone, M., Dancík, V., Wagner, B.K., and Clemons, P.A. (2013). Target identification and mechanism of action in chemical biology and drug discovery. Nat. Chem. Biol. 9, 232–240.

28 Garreau de Loubresse, N., Prokhorova, I., Holtkamp, W., Rodnina, M.V., Yusopova, G., and Yusupov, M. (2014). Structural basis for the inhibition of the eukaryotic ribosome. Nature 513, 517–522.

29 Sandvig, K., and van Deurs, B. (2002) Transport of protein toxins into cells: pathways used by ricin, cholera toxin and Shiga toxin. FEBS Lett. 529, 49–53.

30 Derakhshankhah, H., and Jafari, S. (2018). Cell penetrating peptides: A concise review with emphasis on biomedical applications. Biomed. Pharmacother. 108, 1090–1096.

31 Hong, V., Presolski, S.I., Celia, M., and Finn, M.G. (2009). Analysis and optimization of copper-catalyzed azide-alkyne cycloaddition for bioconjugation. Angew. Chem. Int. Ed. 48, 9879–9883.

32 Meldal, M., and Tornøe, C.W. (2008). Cu-catalyzed azide-alkyne cycloaddition. Chem. Rev. 108, 2952–3015.

33 Hein, J.E., and Fokin, V.V. (2010). Copper-catalyzed azide–alkyne cycloaddition (CuAAC) and beyond: new reactivity of copper(I) acetylides. Chem. Soc. Rev. 39, 1302–1315.

34 Varkouhi, A. K., Scholte, M., Storm, G., Haisma, H. J. (2011). Endosomal escape pathways for delivery of biologicals. J. Control. Release. 151, 220–228.

35 Williamson, N.R., Commander, P.M. B. and Salmond, G.P.C. (2010). Quorum sensing-controlled Evr regulates a conserved cryptic pigment biosynthetic cluster and a novel phenomycin-like locus in the plant pathogen, Pectobacterium carotovorumem. Environ. Microbiol. 12, 1811–1827.

36 Drozdetskiy, A., Cole, C., Procter, J., Barton, G.J. (2015) JPred4: a protein secondary structure prediction server, Nucleic Acids Research, 43, W389–W394

37 Nakamura, S., Takeuchi, T., Hori, S., Matsuzaki, M., and Umezawa, H. (1971). Phenomycin, toxicity and distribution. J. Antibiot. 24, 197–199.

38 Wang, F., Xiao, J., Pan, L., Yang, M., Zhang, G., Jin, S., Yu, J. (2008). A Systematic Survey of Mini-Proteins in Bacteria and Archaea, PLoS ONE, 3, e4027.

39 Conibear, A.C., Chaousis, S., Durek, T., Rosengren, K.J., Craik, D.J., Schroeder, C.I. (2016). Approaches to the stabilization of bioactive epitopes by grafting and peptide cyclization, Biopolym. 106, 89–100.

40 Andersen, K.R., Leksa, N.C., Schwartz, T.U. (2013). Optimized E. coli expression strain LOBSTR eliminates common contaminants from His-tag purification. Proteins, 81, 1857–1861.

41 Ben-Shem, A., Garreau de Loubresse, N., Melnikov, S., Jenner, L.B., Yusupova, G., Yusupov, M. (2011). The structure of the eukaryotic ribosome at 3.0 Å resolution. Science, 334, 1524–1529.

42 Maguire, B.A., Wondrack, L.M., Contillo, L.G., Xu, Z. (2008). A novel chromatography system to isolate active ribosomes from pathogenic bacteria. RNA, 14, 188–195.

43 Nielsen, J.T., Kulminskaya, N., Bjerring, M., Nielsen, N.C. (2014). Automated robust and accurate assignment of protein resonances for solid state NMR. J. Biomol. NMR, 59, 119–134.

44 Shen, Y., Bax, A. (2015). Protein Structural Information Derived from NMR Chemical Shift with the Neural Network Program *TALOS-N*, Methods Mol. Biol. 1260, 17–32.

45 Güntert P. (2004) Automated NMR Structure Calculation With CYANA. In: Downing A.K. (eds) Protein NMR Techniques. Methods in Molecular Biology, vol 278. Humana Press.

46 Frishman, D., Argos, P. (1995). Knowledge-based protein secondary structure assignment, Proteins, 23, 566–579.

47 Bhattacharya, A., Tejero, R., Montelione, G.T. (2007). Evaluating protein structures determined by structural genomics consortia, Proteins, 66, 778–795.

