## Supplementary Information for "Structure and function of the bacterial protein toxin phenomycin"

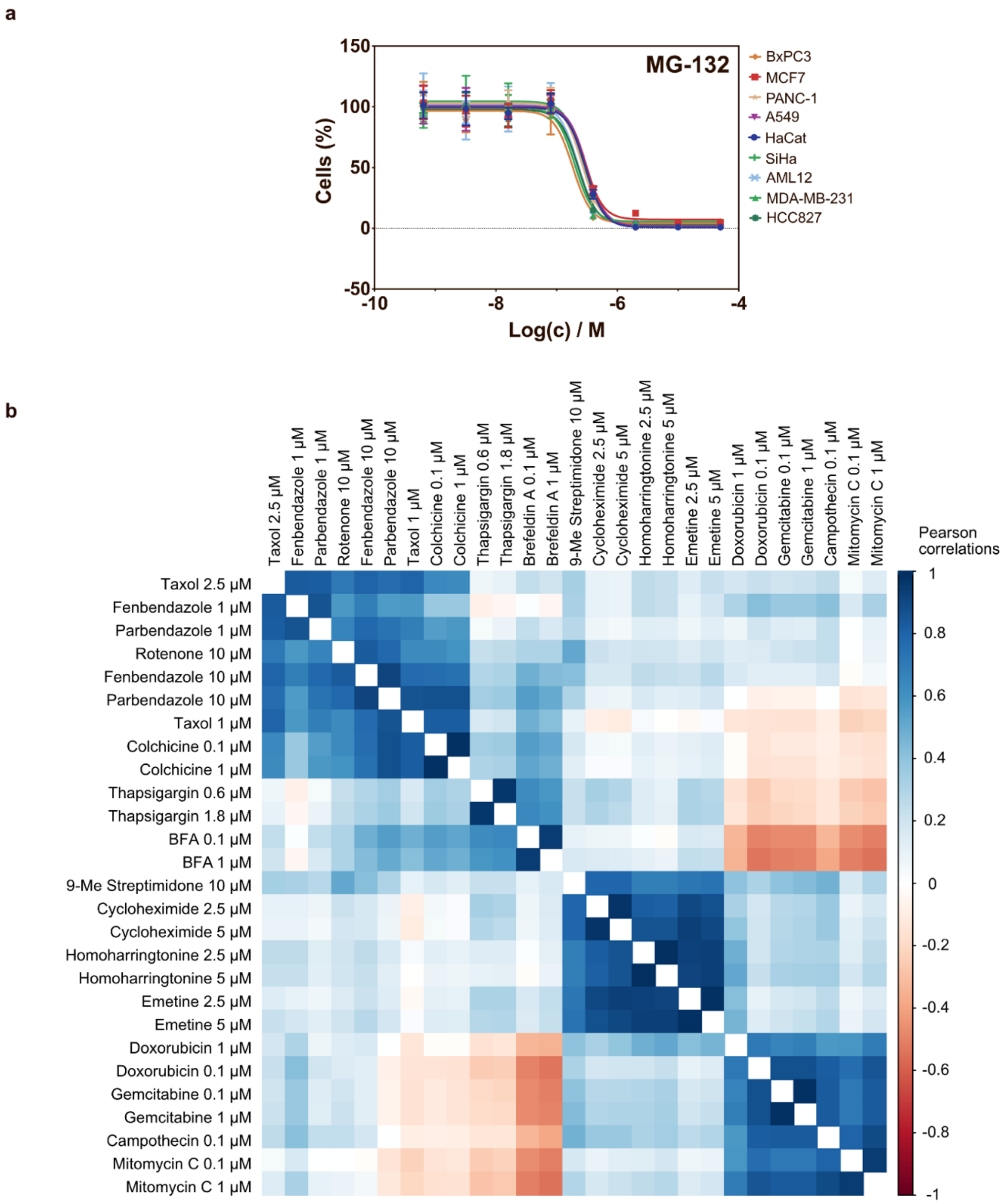


**Figure S1. Supplementary viability and morphological profiling data.** (a) Viability of several cell lines after treatment with MG-132 for 72 h. All data points represent mean ± s.d. (*N*=3). (b) Correlation matrix of morphological profiles of various biologically active compounds. Compounds with similar mechanism of action forms distinct clusters: Tubulin modulators (taxol, rotenone, parbendazole, fenbendazole, colchicine), ER stress inducers (brefeldin A, thapsigargin), translation inhibitors (9-Me streptimidone, cycloheximide, emetine, homoharringtonine) and DNA interferers (doxorubicin, gemcitabine, camptothecin, mitomycin C). For the data shown here all compounds are dosed as 25 µL 4X in 2 % DMSO with no addition of PBS. In the data in figure 1 and S6 peptides are supplied in 13% PBS with addition of DMSO to a final concentration of 0.5 % DMSO and 3.25 % PBS in the wells. This difference has a slight effect on the profiles and thus these two situations are kept separate. Related to Figure 1.

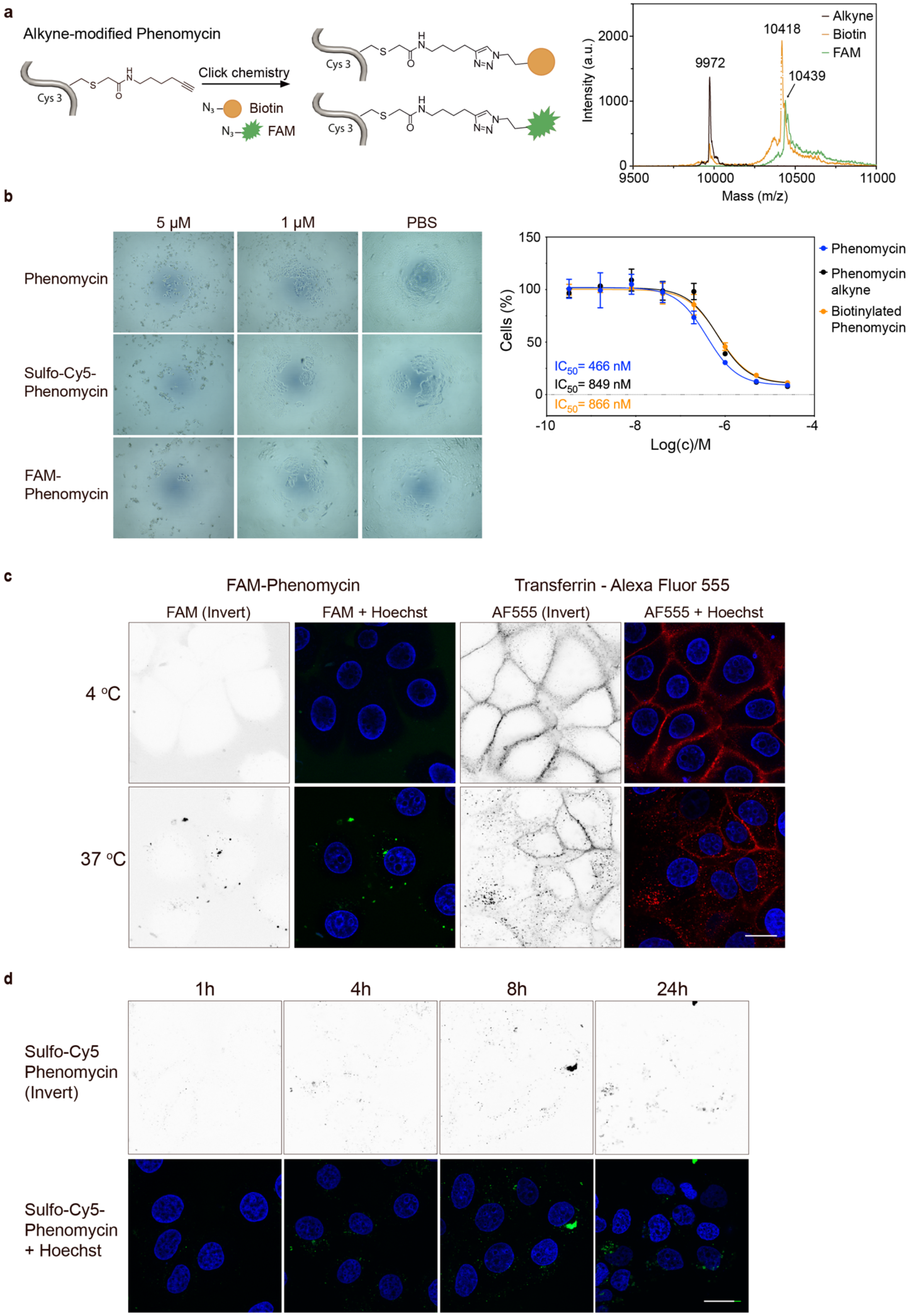

a
Alkyne-modified Phenomycin
Cys 3
Click chemistry
Biotin
FAM
Intensity (a.u.)
Mass (m/z)
9972
10418
10439
Alkyne
Biotin
FAM
b
5 μM
1 μM
PBS
Phenomycin
Sulfo-Cy5-Phenomycin
FAM-Phenomycin
Cells (%)
Log(c)/M
IC50= 466 nM
IC50= 849 nM
IC50= 866 nM
Phenomycin
Phenomycin alkyne
Biotinylated Phenomycin
c
FAM-Phenomycin
Transferrin - Alexa Fluor 555
FAM (Invert)
FAM + Hoechst
AF555 (Invert)
AF555 + Hoechst
4 °C
37 °C
d
1h
4h
8h
24h
Sulfo-Cy5 Phenomycin (Invert)
Sulfo-Cy5-Phenomycin + Hoechst

**Figure S2. Modified phenomycin maintains toxicity in mammalian cells and its cellular uptake is enhanced over time but can be impaired at low temperature.** (a) Alkyne-modified phenomycin is conjugated to FAM-azide and biotin-azide by copper-catalyzed azide-alkyne cycloaddition followed by preparative reverse phase HPLC purification and MALDI measurements. MALDI-data (uncalibrated) of the different conjugates are shown. Calculated masses: Alkyne: 9963.16, Biotin: 10407.71 FAM: 10421.16. (b) Viability of cells after treatment with modified phenomycins for 72 h. MCF7 cells were treated with phenomycin and fluorophore-labelled phenomycins (1 µM and 5 µM) for 72 h and cells were imaged by light microscopy. For alkyne- and biotin-modified phenomycins, cell viability was assessed with the CellTiter-Blue assay. (c) At low temperature, the cellular uptake of FAM-phenomycin was blocked similar to transferrin. MCF7 cells were treated with FAM-phenomycin (20 µM) or transferrin Alexa fluor 555 conjugate (25 µg/ml) for 4 h at 4 $^{o}$C or 37 $^{o}$C and imaged using confocal microscopy. Cell nuclei are stained by Hoechst 33342 (blue). Scale bar is 20 µm. 8-bit images, display range: 0-150 (d) Time course study of sulfo-Cy5-phenomycin (1 µM) uptake in MCF7 cells. Scale bar is 20 µm. 8-bit images, display range: 0-120 (Hoechst). Related to Figure 2.

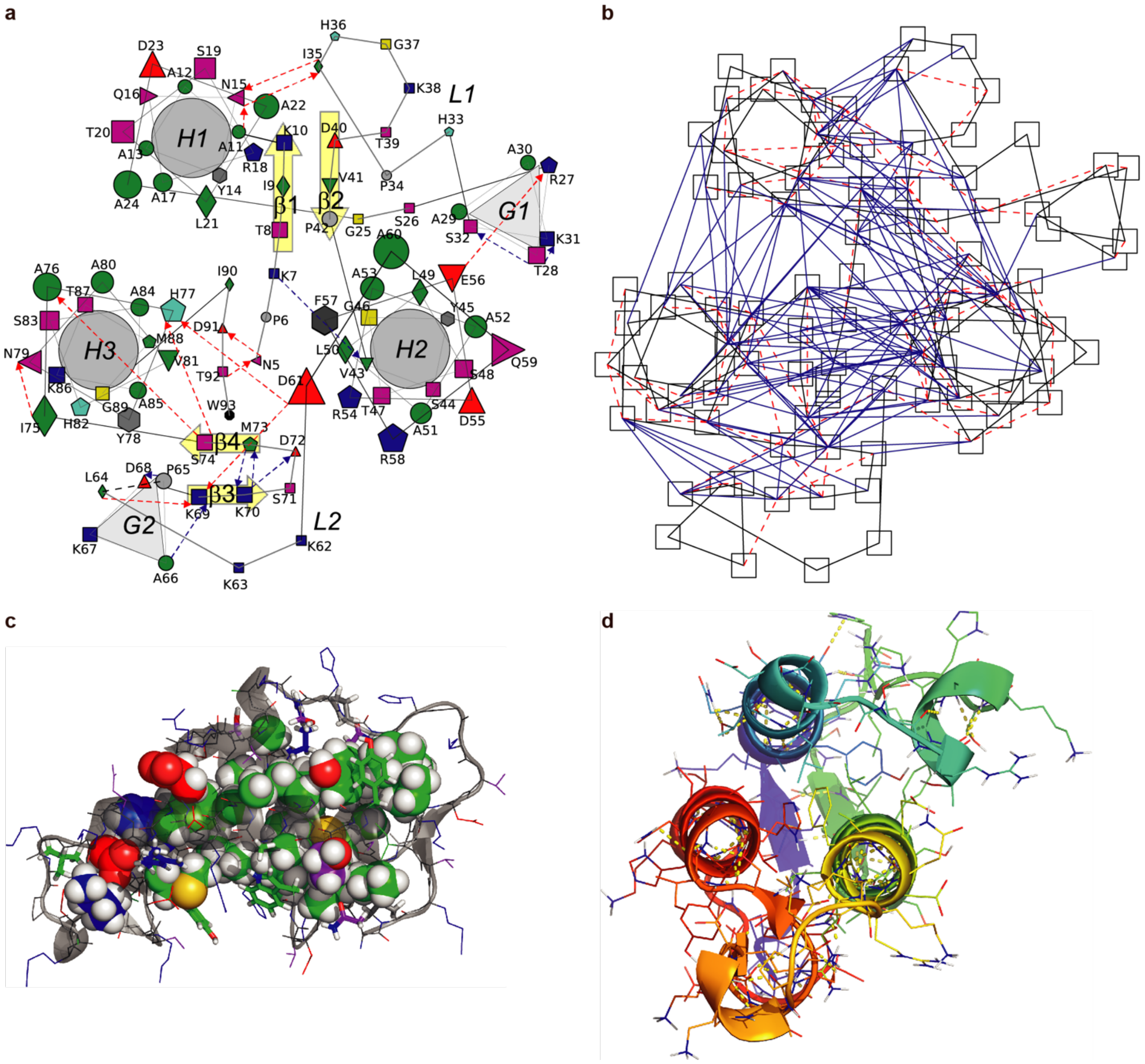


**Figure S3. Supplementary illustrations of the structure of phenomycin.** (a) Schematic illustration based on a view down the helix axes showing helices as helical wheel projections and placing exposed residues nearest the edges of the illustration. The three alpha-helices H1 (A11-A24), H2 (V43-D61) and H3 (I75-M88) are labeled and highlighted with grey circles and the $3_{10}$-helixes are labeled: G1 (A29-K31) and G2 (A66-D68) and highlighted with light grey triangles, and beta-strands β1-4 (T8-K10, D40-P42, K69-K70, M73-S74) are labelled and highlighted with yellow filled arrows. Loops 1 and 2 are indicated with L1 and L2. Each amino acid type is shown by a symbol colored as in Fig. 3C but using specific aromatic(grey), His(cyan), marked with: A(circles), STGK(squares), YF(hexagons), W(octagon), IL(diamonds), V(triangle pointing up), RHM(pentagons), and D/E/N/Q by triangles pointing up/down/left/right, respectively. Non-$\alpha$-helical main- chain hydrogen bonds are indicated with blue dashed arrows connecting the symbols from O-atom acceptor to $H_N$-atom donor whereas side chain H-bonds are denoted using red dashed arrows connecting symbols corresponding to from acceptor to donor. (b) Visualization of distribution of non-trivial distances constraints (see Table S3) between residues showing long-range (residues number difference, Δ > 4) and medium-range (Δ = 3 or Δ = 4) restraints with blue lines and red dashed lines, respectively, and showing each residue as a square positioned as in (a) connected with black lines. (c) Structure representation of hydrophobic core residues (see also footnote c to Table S3) showing buried residues, (relative solvent accessibility, RSA < 0.22) with spheres for all atoms, semi buried/exposed residues (0.35 < RSA < 0.22) with sticks, and exposed residues (RSA > 0.35) using lines for heavy atoms only and using colors as in Figure 3d throughout. (d) Rainbow cartoon representation of Phenomycin as in Figure 3a viewed down the helix axes showing also heavy atoms and polar hydrogens of the side chains colored as in Figure 3d and hydrogen bonds using yellow dashes. Related to Figure 3.

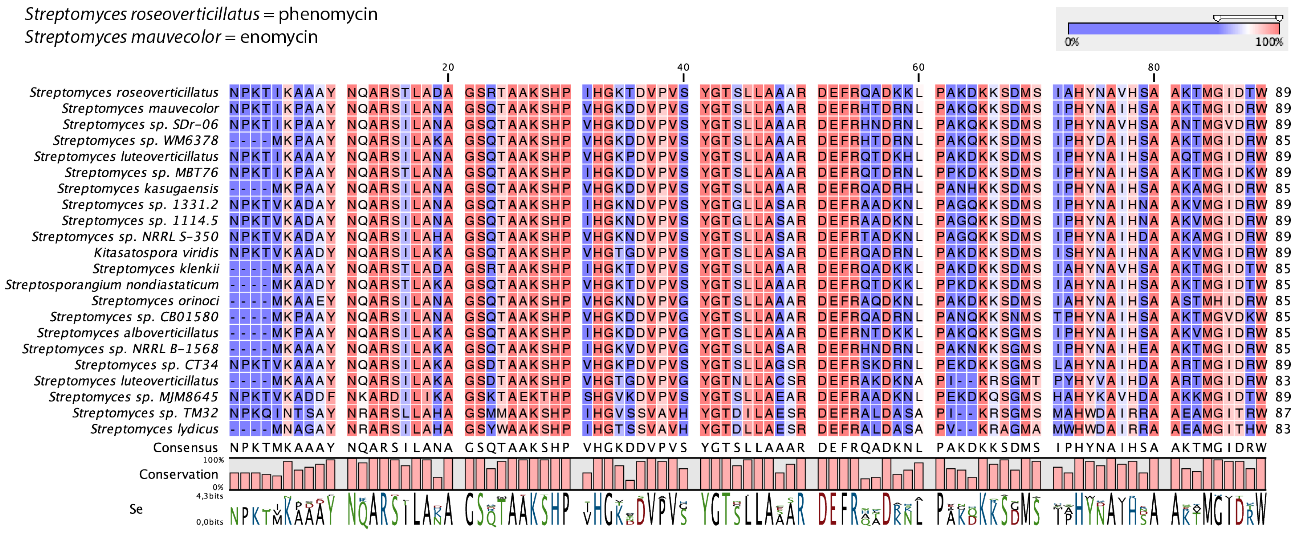

**Figure S4. Protein alignment of phenomycin and enomycin with predicted proteins from other Streptomyces and Kitasatospora.** The initial blast search for similar proteins gave a list of several sequences that all shared a high sequence similarity with phenomycin and enomycin. The result of a multiple alignment made using CLC Main Workbench with default settings is shown above. The blue/red color scale indicates an overall high similarity with more variance in the region constituting loop 2. Related to Figure 4.

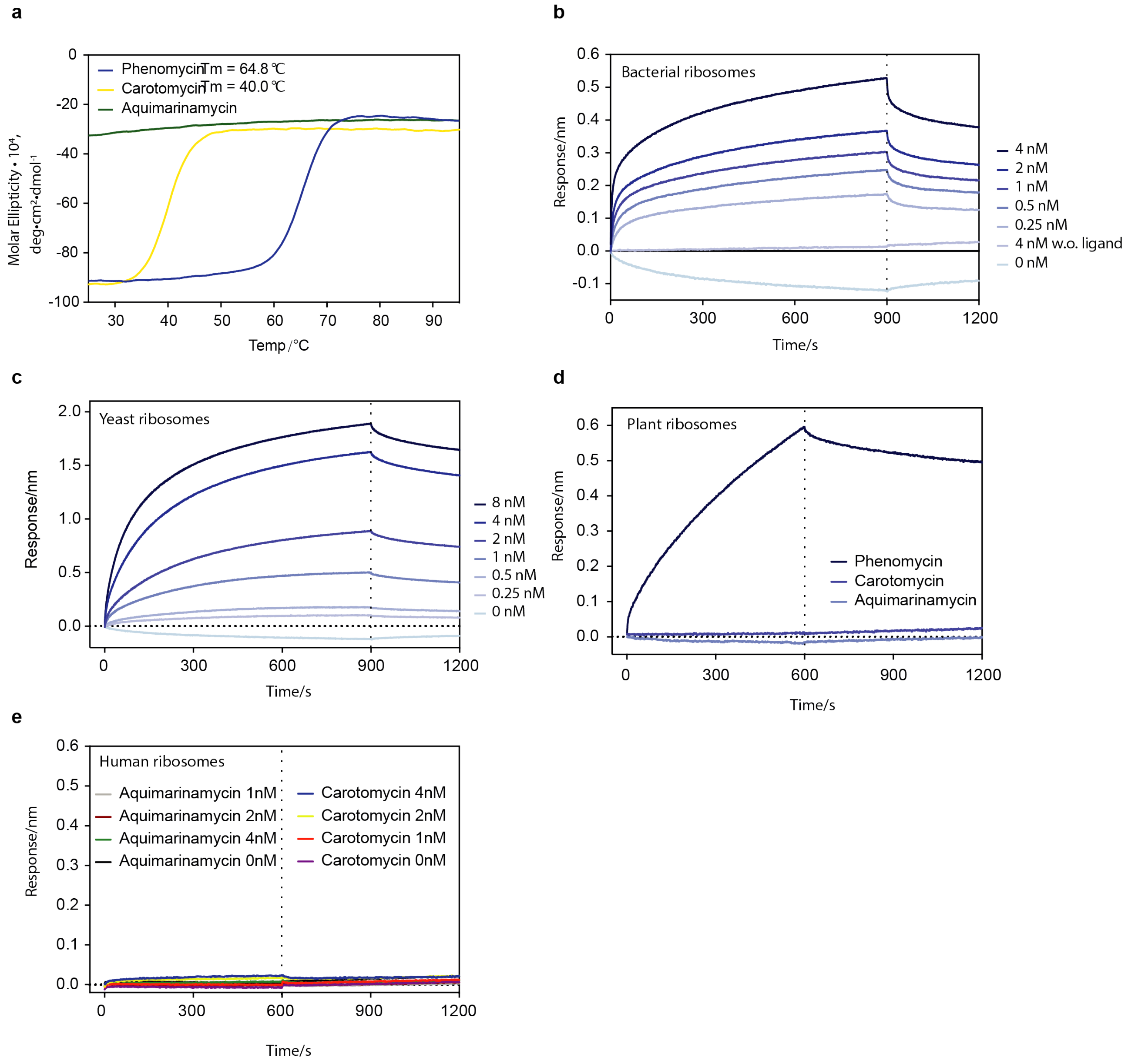


**Figure S5. Stability and binding of phenomycin and homologs.** (a) Temperature stability of phenomycin (blue), carotamycin (yellow), and aquamarinamycin (green) measured by CD temperature scan at 222 nm. Melting temperatures (Tm) are indicated on graph. (b-c) Binding of bacterial or yeast ribosomes to phenomycin immobilized on streptavidin BLI sensors. (d) Binding of plant ribosomes to phenomycin, carotamycin and aquimarinamycin. (e) Binding studies of human ribosomes to carotamycin and aquamarinamycin immobilized on streptavidin BLI sensors. Related to figure 5.

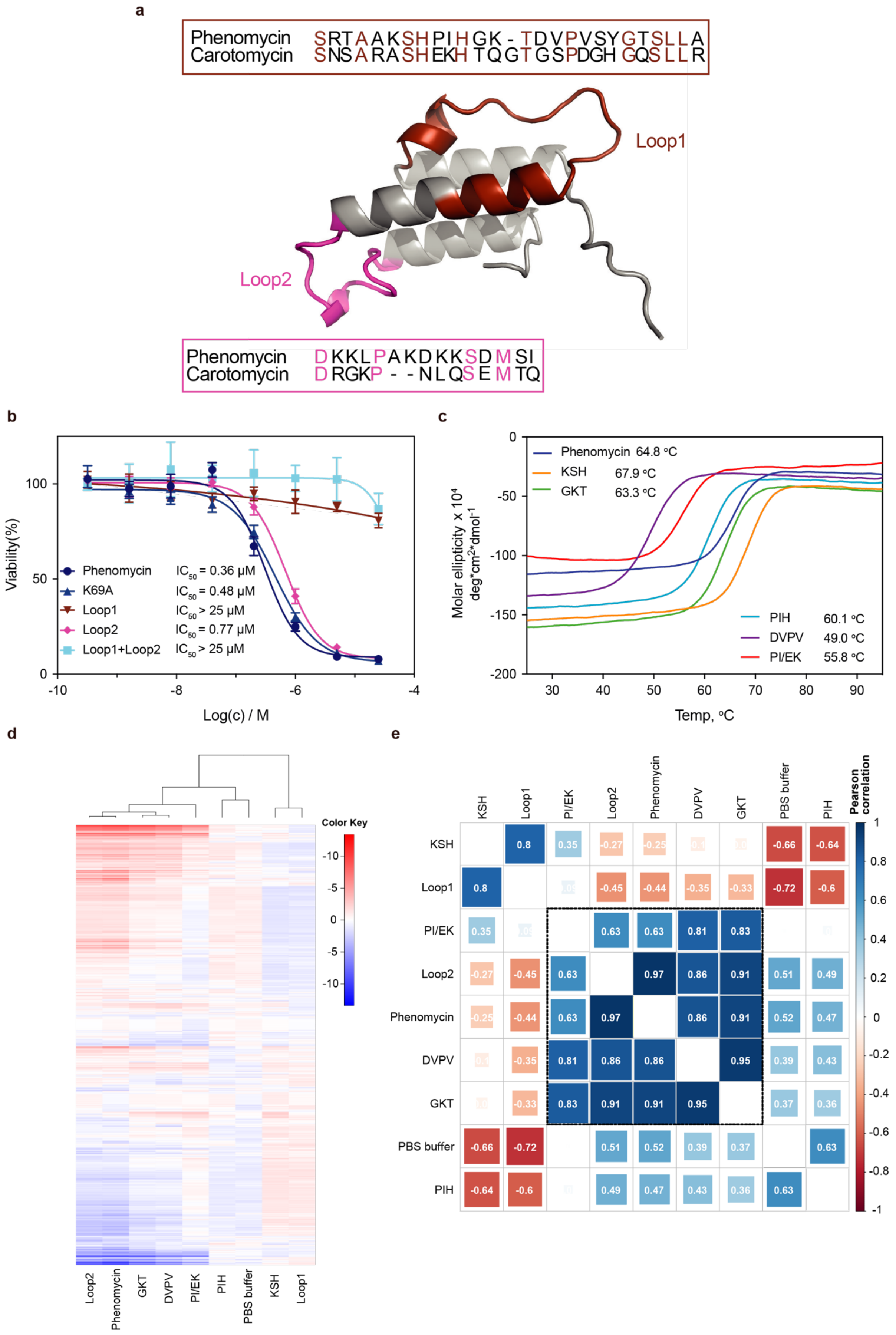

a
Phenomycin SRTAAKSHPIHGK-TDVPVSYGTSLLA
Carotomycin SNSARASHEKHTQGTGSPDGHGQSLLR
Loop1
Loop2
Phenomycin DKKLPAKDKKSDMSI
Carotomycin DRGKP--NLQSEMTQ
b
Viability(%)
Log(c) / M
Phenomycin IC50 = 0.36 μM
K69A IC50 = 0.48 μM
Loop1 IC50 > 25 μM
Loop2 IC50 = 0.77 μM
Loop1+Loop2 IC50 > 25 μM
c
Molar ellipticity x 10^4 deg*cm^2*dmol^-1
Temp, °C
Phenomycin 64.8 °C
KSH 67.9 °C
GKT 63.3 °C
PIH 60.1 °C
DVPV 49.0 °C
PI/EK 55.8 °C
d
Color Key
Loop2
Phenomycin
GKT
DVPV
PI/EK
PIH
PBS buffer
KSH
Loop1
e
KSH
Loop1
PI/EK
Loop2
Phenomycin
DVPV
GKT
PBS buffer
PIH
Pearson correlation

**Figure S6. Detailed investigations of the two loop regions found in phenomycin.** (a) The three-dimensional structure of phenomycin highlights the two loop regions found. To assess the functional implications of changing sequence in the loops we exchanged the phenomycin sequence with the corresponding sequence found in carotomycin. (b) MCF7 cell viability after treatment with phenomycin and four generated analogs. K69A is a single point mutation generated because K69 appeared to play a key structural role in loop 1 as revealed by the NMR structure. Phenomycin with Loop2 substituted with sequence from carotomycin (Loop2, light blue) has no effect on bioactivity while phenomycin with Loop1 from carotomycin (Loop1, red) or a combination of both significantly reduce bioactivity demonstrating the importance of Loop1. K69A has no effect on bioactivity. (c) Temperature scans using CD to compare the stability of the secondary structure when introducing three/four amino acid substitutions (to alanines) in segments of loop 1. (d) Morphological profiles of phenomycin mutants (10 µM) and negative control (phosphate buffered saline buffer (PBS buffer)). U2OS cells were treated for 24 hours in four replicates which were averaged to one fingerprint. Profiles are clustered by hierarchical clustering with average distance and Pearson correlation as the distance metric. (e) Pearson correlation matrix of profiles in (d). Related to Figure 6.

**Table S1. Overview of different proteins used in the study. Related to Figure 1-6 and Figure S1-S6.**

| | **Protein sequence (the cleaved protein is written in bold, the N-terminal amino acids that differ from the naturally occurring protein is in italic, the specific changes relative to the wt proteins are in red)** |
|---|---|
| Carotomycin | MKHHHHHHPMSDYDIPTTENLYFQ***GA*MAISDNDYNRARAILIAAGSNSARASHEKHTQGTGSPDGHGQSLLRGSRDEFRTADRGKPNLQSEMTQVHYNAIHAAAQQMGIPNW** |
| M3C carotomycin | MKHHHHHHPMSDYDIPTTENLYFQ***GAC*AISDNDYNRARAILIAAGSNSARASHEKHTQGTGSPDGHGQSLLRGSRDEFRTADRGKPNLQSEMTQVHYNAIHAAAQQMGIPNW** |
| Carotomycin Loop1 | MKHHHHHHPMSDYDIPTTENLYFQ***GA*MAISDNDYNRARAILIAAGSRTAAKSHPIHGKTDVPVSYGTSLLAGSRDEFRTADRGKPNLQSEMTQVHYNAIHAAAQQMGIPNW** |
| Carotomycin Loop2 | MKHHHHHHPMSDYDIPTTENLYFQ***GA*MAISDNDYNRARAILIAAGSNSARASHEKHTQGTGSPDGHGQSLLRGSRDEFRTADKKLPAKDKKSDMSIVHYNAIHAAAQQMGIPNW** |
| Carotomycin Loop1+Loop2 | MKHHHHHHPMSDYDIPTTENLYFQ***GA*MAISDNDYNRARAILIAAGSRTAAKSHPIHGKTDVPVSYGTSLLAGSRDEFRTADKKLPAKDKKSDMSIVHYNAIHAAAQQMGIPNW** |
| Aquimarinamycin | MKHHHHHHPMSDYDIPTTENLYFQ***GA*MGPIKDRDYNKAKALLEAAGSKSAEKSHPKHTGSKGSPDSHGRSLYNEAQQDFGGQPTAAQLEALRKASRFLGV** |
| M3C aquimarinamycin | MKHHHHHHPMSDYDIPTTENLYFQ***GAC*GPIKDRDYNKAKALLEAAGSKSAEKSHPKHTGSKGSPDSHGRSLYNEAQQDFGGQPTAAQLEALRKASRFLGV** |
| Phenomycin | MKHHHHHHPMSDYDIPTTENLYFQ***GAMA*NPKTIKAAAYNQARSTLADAGSRTAAKSHPIHGKTDVPVSYGTSLLAAARDEFRQADKKLPAKDKKSDMSIAHYNAVHSAAKTMGIDTW** |
| M3C phenomycin | MKHHHHHHPMSDYDIPTTENLYFQ***GACA*NPKTIKAAAYNQARSTLADAGSRTAAKSHPIHGKTDVPVSYGTSLLAAARDEFRQADKKLPAKDKKSDMSIAHYNAVHSAAKTMGIDTW** |
| K69A | MKHHHHHHPMSDYDIPTTENLYFQ***GAMA*NPKTIKAAAYNQARSTLADAGSRTAAKSHPIHGKTDVPVSYGTSLLAAARDEFRQADKKLPAKDAKSDMSIAHYNAVHSAAKTMGIDTW** |
| Phenomycin Loop1 | MKHHHHHHPMSDYDIPTTENLYFQ***GAMA*NPKTIKAAAYNQARSTLADAGSNSARASHEKHTQGTGSPDGHGQSLLRAARDEFRQADKKLPAKDKKSDMSIAHYNAVHSAAKTMGIDTW** |
| Phenomycin Loop2 | MKHHHHHHPMSDYDIPTTENLYFQ***GAMA*NPKTIKAAAYNQARSTLADAGSRTAAKSHPIHGKTDVPVSYGTSLLAAARDEFRQADRGKPNLQSEMTQAHYNAVHSAAKTMGIDTW** |
| Phenomycin Loop1+Loop2 | MKHHHHHHPMSDYDIPTTENLYFQ***GAMA*NPKTIKAAAYNQARSTLADAGSNSARASHEKHTQGTGSPDGHGQSLLRAARDEFRQADRGKPNLQSEMTQAHYNAVHSAAKTMGIDTW** |
| KSH-AAA | MKHHHHHHPMSDYDIPTTENLYFQ***GAMA*NPKTIKAAAYNQARSTLADAGSRTAAAAAPIHGKTDVPVSYGTSLLAAARDEFRQADKKLPAKDKKSDMSIAHYNAVHSAAKTMGIDTW** |
| PIH-AAA | MKHHHHHHPMSDYDIPTTENLYFQ***GAMA*NPKTIKAAAYNQARSTLADAGSRTAAKSHAAAGKTDVPVSYGTSLLAAARDEFRQADKKLPAKDKKSDMSIAHYNAVHSAAKTMGIDTW** |
| GKT-AAA | MKHHHHHHPMSDYDIPTTENLYFQ***GAMA*NPKTIKAAAYNQARSTLADAGSRTAAKSHPIHAAADVPVSYGTSLLAAARDEFRQADKKLPAKDKKSDMSIAHYNAVHSAAKTMGIDTW** |
| DVPV-AAAA | MKHHHHHHPMSDYDIPTTENLYFQ***GAMA*NPKTIKAAAYNQARSTLADAGSRTAAKSHPIHGKTAAAASYGTSLLAAARDEFRQADKKLPAKDKKSDMSIAHYNAVHSAAKTMGIDTW** |
| PI-EK | MKHHHHHHPMSDYDIPTTENLYFQ***GAMA*NPKTIKAAAYNQARSTLADAGSRTAAKSHEKHGKTDVPVSYGTSLLAAARDEFRQADKKLPAKDKKSDMSIAHYNAVHSAAKTMGIDTW** |

**Table S2: Imaging settings for morphological profiling. Related to Figure 1 and Figure S6.**

| Channel | Dyes | Excitation LED | Beamsplitter | Emission filter |
|---|---|---|---|---|
| DNA | Hoechst 33342 | 385 nm | RTBS 405 + 493 + 610 | TBP 425/30 + 524/50 + 688/154 |
| ER | Concanavalin-AF488 | 470 nm | RTBS 405 + 493 + 610 | TBP425/30 + 524/50 + 688/145 |
| RNA | SYTO 14 green fluorescent nucleic acid stain | 511 nm | RTBS 450 + 538 + 610 | TBP 467/24 + 555/25 + 687/145 |
| AGP | Phalloidin-AF568 + Wheat-germ agglutinin-AF555 | 567 nm | RQBS 405 + 493 + 575 + 653 | QBP 425/30 + 514/30 + 592/25 + 709/100 |
| Mito | MitoTracker Deep Red | 625 nm | RQBS 405 + 493 + 575 + 653 | QBP 425/30 + 514/30 + 592/25 + 709/100 |

**Table S3. Restraint and structure statistics for phenomycin. Related to Figure 3 and Figure S3.**

| | |
|---|---|
| **Restraint statistics** | |
| Distance Restraints [a] | |
| Total | 1901 |
| Intra-residue | 408 |
| Sequential | 519 |
| Medium range | 431 |
| Long range | 543 |
| Dihedral angle | 150 |
| Restraint violations: number of / rms (max) | |
| Distance violations > 0.1 Å | 2.9/0.0048 (0.16) |
| Distance violations > 0.2 Å | 0.3/0.0048 (0.16) |
| Torsion violations > 5 ° | 0.0/0.684 (3.05) |
| **Structure statistics** | |
| Coordinate r.m.s.d. to mean: backbone/heavy (Å)[b] | |
| structured (res. 5-93) | 0.30/0.76 |
| all | 0.47/0.83 |
| buried core [c] | 0.21/0.46 |
| Angle rms deviation from ideal (°) | 0.2 |
| Bond rms deviation from ideal (Å) | 0.001 |
| Ramachandran statistics [d] | |
| Most favoured regions | 93.2% |
| Allowed regions | 6.6% |
| Disallowed regions | 0.2% |
| Structure Quality Validation Metrics [e] (raw / Z-score) | |
| Procheck G-factor (phi / psi only) | -0.02/0.24 |
| Procheck G-factor (all dihedral angles) | -0.38/-2.25 |
| Verify3D | 0.32/-2.25 |
| ProsaII (-ve) | 0.75/0.41 |
| MolProbity clashscore | 30.14/-3.65 |

[a] Long-range meaning that the residue difference, D, was 5 or more and medium range; 1<D<5.

[b] Coordinate rmsd calculated for 20 ensemble members

[c] Residues with a relative accessible surface area (RSA) > 0.23: residues 9, 11, 13-14, 17, 21, 29, 35, 41, 43, 46-47, 49-50, 52-53, 56-57, 61, 69, 73-74, 77, 80-81, 84-85, 88, and 90. The accessible surface area was calculated using DSSP #REF# normalizing with theoretical maximal values by Tien et al. #REF: doi:10.1371/journal.pone.0080635#

[d] MolProbity Ramachandran statistics

[e] Calculated using the Protein Structure Validation Suite (http://psvs-1_5-dev.nesg.org). A positive Z-score indicates a "better" score.
